# Flux modelling analysis reveals the metabolic impact of cryptic plasmids and environmental conditions in probiotic *Escherichia coli* Nissle 1917

**DOI:** 10.1101/2025.11.17.688048

**Authors:** Paola Corbín-Agustí, Alba Arévalo-Lalanne, Patricia Álvarez, Maria Enrique, Daniel Ramón, Juli Peretó, Marta Tortajada

## Abstract

*Escherichia coli* Nissle 1917 (EcN) is a well-characterized Gram-negative probiotic distinguished by its unique, strain-specific physiology. Genome-scale metabolic models (GEMs) are powerful tools for elucidating metabolic traits and predicting genotype–phenotype relationships. Although several EcN GEMs have been published, none have explicitly represented its probiotic physiology. Here, we present a manually curated GEM of EcN that, for the first time, incorporates the energetic costs associated with its cryptic plasmids. Inclusion of a plasmid-specific module improved biomass yield predictions and overall model accuracy, providing a more physiologically realistic representation of EcN metabolism. Using COBRA methodologies and possibilistic metabolic flux analysis, this model and previous EcN reconstructions were systematically compared to evaluate the trade-off between model complexity and predictive performance. The analysis revealed that increased structural detail does not necessarily enhance quantitative accuracy and that predictive reliability depends on both computational methodology and model context. Metabolomic profiling under gut-like anaerobic conditions further showed that EcN exhibits a distinctive metabolic phenotype, characterized by elevated amino acid consumption and enhanced short-chain fatty acid production. These findings highlight the unique probiotic physiology of EcN and demonstrate the utility of metabolic modeling for reproducing and exploring such traits. Overall, this study provides a quantitatively reliable and physiologically relevant framework for modeling *E. coli* Nissle 1917 and related commensal bacteria, supporting advances in probiotic engineering, synthetic biology, and bioprocess design.

**Graphical Abstract Summary:** This study presents a manually curated genome-scale model of *Escherichia coli* Nissle 1917 that accounts for the metabolic cost of its cryptic plasmids. Through systematic comparison with previous reconstructions and validation against fluxomics datasets, the models improved accuracy in predicting growth and fluxes. Simulations and experiments under gut-like conditions provide new insights into EcN’s unique probiotic traits.

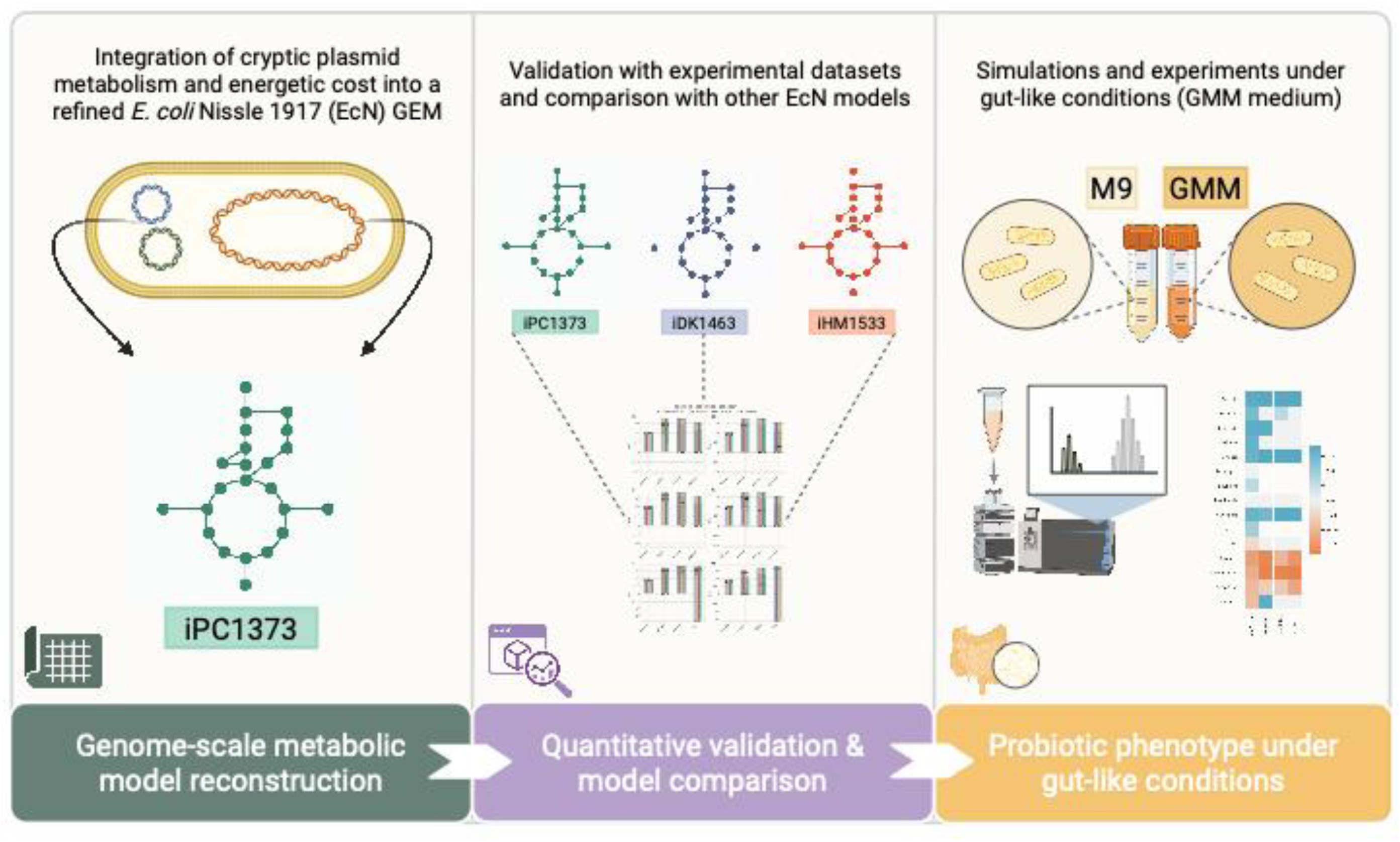

## 1. Introduction

*Escherichia coli* Nissle 1917 (EcN) is a well-characterized probiotic strain originally isolated by Alfred Nissle in 1917 from the faeces of a First World War soldier who appeared resistant to dysentery (Beimfohr, 2016). Since its discovery, EcN has been extensively used to treat gastrointestinal disorders such as infectious diarrhea, inflammatory bowel diseases (IBD), ulcerative colitis or Crohn’s disease (Petersen *et al*., 2011; Olier *et al*., 2012; Maltby *et al*., 2013). Remarkably, EcN remains the only Gram-negative strain in routine probiotic use, despite belonging to a species typically associated with pathogenic lineages.

EcN exhibits several unique, strain-specific features that underpin its probiotic efficacy. These include (i) high fitness in the intestinal environment and stable colonization through strong adhesion to the epithelium, biofilm formation, and antimicrobial resistance; (ii) competitive exclusion of enteric pathogens via production of antimicrobials such as microcins, interference with virulence mechanisms, and sequestration of key nutrients such as iron; and (iii) reinforcement of host mucosal immunity by activating TLR and NF-κB signaling, enhancing epithelial barrier function, and stimulating antimicrobial peptide production (Deriu *et al*., 2013; Schumann *et al*., 2014; Hare *et al*., 2022). EcN also lacks typical virulence factors such as α-hemolysin and fimbriae and displays a semi-rough lipopolysaccharide, which may contribute to immune evasion, attenuated inflammatory response, and improved mucosal adhesion (Deriu *et al*., 2013; Schumann *et al*., 2014; Hare *et al*., 2022).

Genome-scale metabolic models (GEMs) represent the metabolic capabilities of an organism through the stoichiometric formulation of its biochemical reactions. Over the past two decades, GEMs have become indispensable powerful and robust computational tools in systems biology for elucidating microbial metabolism, predicting genotype–phenotype relationships, functional characteristics, and optimizing industrial bioprocesses (Monk *et al*., 2013, 2016, 2017; Pearcy *et al*., 2021; Bernstein *et al*., 2023). Due to the large number of reactions and metabolic uncertainties, GEMs are inherently underdetermined and require constraint-based approaches—collectively known as COBRA (Constraint-Based Reconstruction and Analysis)—to predict metabolic flux distributions. Among the most widely applied methods are metabolic flux analysis (MFA), which integrates experimental flux measurements to estimate flux rates, and flux balance analysis (FBA), which uses stoichiometric constraints and optimization principles to infer metabolic states under an assumed cellular objective (Orth *et al.,* 2010; Antoniewicz, 2021). Complementary to these approaches, possibilistic MFA provides a framework for handling uncertainty by combining constraint-based models with measurement error distributions represented as possibility functions. This method enables robust flux estimation even when experimental data are incomplete, using linear programming to identify feasible flux ranges (Llaneras *et al*., 2009; Llaneras, 2011).

Recent GEM-based studies have explored the metabolic capabilities of EcN. These models have been used to benchmark EcN against laboratory or pathogenic *E. coli* strains, identify distinctive metabolic features and explore its potential as a chassis for biosynthetic applications (Kim *et al*., 2021; Moutinho *et al*., 2022; van’t Hof *et al*., 2022).

EcN harbors two cryptic plasmids, pMUT1 (3.2 kb) and pMUT2 (5.5 kb), both possessing ColE1-type replication origins and maintained at high copy numbers (more than 100 copies per cell) (Zainuddin *et al*., 2019). These plasmids are stably inherited and commonly used for strain typing (Blum-Oehler *et al*., 2003). Although historically considered phenotypically neutral, recent transcriptomic evidence indicates that deletion of pMUT1 and pMUT2 substantially alters the expression of genes involved in cell aggregation and amino acid metabolism, suggesting possible contributions to EcN’s probiotic function (Lin *et al*., 2024). Additionally, their stable maintenance and replication machinery have been exploited as antibiotic-free expression systems for targeted intestinal delivery of heterologous proteins (Zhou *et al*., 2024).

However, plasmid carriage often imposes a significant metabolic burden on host cells due to the energetic costs of replication and gene expression. This phenomenon is well documented in *E. coli*, particularly in recombinant protein production systems (Rozkov *et al*., 2004; Zeng and Yang, 2019). The additional biosynthetic demand typically leads to reduced growth rate and yield compared with plasmid-free strains and is frequently accompanied by metabolic overflow, such as acetate accumulation. Even strains harboring empty, high-copy-number ColE1-derived plasmids exhibit globally altered physiological states and increased non-growth-associated maintenance energy (Ow *et al*., 2009; Oftadeh and Hatzimanikatis, 2024). Several modeling studies have successfully incorporated plasmid-related metabolic costs into constraint-based metabolic models, including GEMs, improving their ability to reproduce experimental data (Özkan *et al*., 2005; Ow *et al*., 2009; Zeng and Yang, 2019; Oftadeh and Hatzimanikatis, 2024). Nevertheless, the explicit consideration of plasmid burden in probiotic EcN models remains limited.

This study presents a manually curated genome-scale metabolic model (GEM) of EcN that explicitly accounts for the energetic costs of its cryptic plasmids. Using COBRA methods and possibilistic MFA, this model and previously published EcN GEMs were systematically compared and evaluated, including prediction of biomass formation and by-product external rates, as well as their consistency with experimental metabolomics and ^13^C-fluxomics datasets. This comparative framework enabled the assessment of how different models predict metabolic behavior under specific experimental and methodological contexts. Importantly, the experimental metabolomics data used were obtained under conditions that mimic those of the probiotic’s native environment, an approach that, to our knowledge, has not been previously applied to this strain. The results provide a mechanistic framework for better understanding the metabolic determinants of EcN’s probiotic properties, and a more accurate characterization of its probiotic capacity and model predictions within that physiological context.

## 2. Materials and Methods

### Bacterial strains and culture conditions

Two strains of *Escherichia coli* were used in this work: *E. coli* Nissle 1917 (DSM 6601) and *E. coli* K12-HB101 (DSM 1607). EcN strain was isolated from Mutaflor commercial product (Ardeypharm) and grown in Lysogeny Broth (LB) at 37°C unless stated otherwise. EcN strain was routinely identified by PCR amplification in cryptic plasmid pMUT2, as described in (Blum-Oehler *et al*., 2003). *E. coli* K12-HB101 strain was purchased from Promega (Madison, WI, USA). Growth media used throughout experiments include M9 minimal medium, with glucose (Monk *et al*., 2016a) and supplemented with casamino acids to fulfill the auxotrophic requirements of K12 - HB101 strain, and rich-gut microbiota medium (GMM), prepared according to Goodman *et al*. (2011) and Średnicka *et al*. (2023) without agar. Composition details are provided in the Supplementary Methods.

### Draft model construction and refinement

The genome sequence of EcN strain was retrieved from NCBI (RefSeq accession number: NZ_CP007799.1; annotated Feb 23, 2017) and used to compare with other reference *E. coli* strains. For that, the protein sequences of K-12 MG1655 (NC_000913) and CFT073 (NC_004431.1) were used to identify the ortholog genes between EcN and these strains by a reciprocal blast best hit method, with an e-value > 1×10^6^ and a percentage of homology greater than 85%. In parallel, the curated models of E. coli K-12 MG1655 (iAF1260, iJO1366 and iML1515) and E. coli CFT073 (ic_1306) were downloaded from the BIGG database (King *et al*., 2016) in JSON format. A first EcN draft model was reconstructed incorporating all shared reactions and metabolites from iAF1260 and ic_1306 metabolic models, dependent on EcN homologous genes. Subsequently, the non-homologous genes of *E. coli* EcN were reviewed and the associated reactions, retrieved from BiGG when possible, were added to the draft model. MetaCyc (Caspi *et al*., 2020), KEGG (Kanehisa *et al*., 2021) and BRENDA (Schomburg *et al*., 2013) databases were also consulted for the manual model curation. The biomass reaction used in this EcN model corresponds to that of the iJO1366 model. Consistency analysis of the topological network was conducted to identify gap metabolites and blocked reactions (Ponce-de-Leon *et al*., 2015). Finally, gaps, reaction directionality and redundancies were manually curated. Reactions, metabolites, genes, and subsystems were annotated using the KEGG, BioCyc, MetaNetX, SEED, Rhea, ChEBI, and NCBI databases. SBOannotator (Leonidou *et al*., 2023) was used to assign SBO terms. The quality of the model was evaluated using the MEMOTE tool (v.0.13.0) (Lieven *et al*., 2020). For the manipulation, study and development of the model Python 3 and the COBRApy library were used (Ebrahim *et al*., 2013). The final GEM is provided in json, mat and sbml format (Supplemental Material S1).

### Metabolic models comparison

The EcN model reconstructed here was compared to the remaining available models (iDK1463 (Kim *et al*., 2021) and iHM1533 (van’t Hof *et al*., 2022) in terms of structure, Memote scores, and predictive analysis.

### Plasmid integration in the EcN model

To incorporate the biosynthesis of the two cryptic plasmids, present in EcN into the model, two reactions were introduced: one representing the consumption of deoxyribonucleotides (dNTPs) and the production of the plasmids (Eq. 1), and a demand reaction (Eq. 2) to enable simulations. The stoichiometric coefficients of dNTPs and pyrophosphate were calculated using the BOFdat algorithm (Lachance *et al*., 2019), based on each plasmid’s nucleotide composition and its estimated copy numbers (Zainuddin *et al*., 2019). The contribution of the plasmids to the cellular dry weight was computed as the DNA fraction, considering the product of plasmids size and copy number relative to the chromosome genome size (Eq. 3).

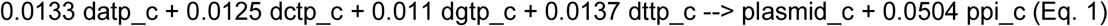

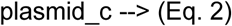

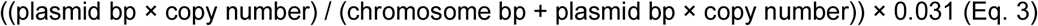

To simulate plasmid replication and its associated metabolic burden, we imposed a fixed lower bound on the plasmid, while keeping biomass maximization as the objective function. To select this fixed value, the strategy was evaluated across a range of NGAM lower bound values, considering which would best reproduce the experimental conditions (see belowe *In silico growth conditions* and *Experimental datasets*) and to assess sensitivity to NGAM. Further details and simulations are provided in the Supplementary Methods.

### ATP expenditure parameters

Growth associated ATP maintenance (GAM) and non-growth associated ATP maintenance (NGAM) are critical parameters for GEMs to accurately represent physiological responses. GAM and NGAM values for wild-type and plasmid-bearing *E. coli* were retrieved from literature. A consensus value of 59.8 mmol·gDW^-1^·h^×1^ for GAM was used (Kim *et al*., 2021). The occurrence of plasmids has been referred to increase NGAM energy due to the metabolic burden associated with their maintenance, replication, and expression (Rozkov *et al*., 2004; Ow *et al*., 2009; Zeng and Yang, 2019). The median of previously reported NGAM values was used to set the lower bound of the ATPM reaction in aerobic (25 mmol·gDW^-1^·h^×1^) and anaerobic conditions (3.5 mmol·gDW^-1^·h^×1^). Further details can be found in Supplementary Methods.

### Experimental datasets employed for model validation

Experimental data were compiled from literature on the growth of EcN on various carbon sources (Revelles *et al*., 2013; Millard *et al*., 2021) and used for model validation. Each dataset contains experimental measurements of several external fluxes, including specific growth rate, carbon source uptake rate, and secretion rates of byproducts. These experimental measurements are provided in Supplementary Methods.

### *In silico* constraints and growth conditions

For reversible reactions, upper and lower bounds of 1000 and -1000 mmol·gDW^-1^·h^×1^, were respectively applied. For irreversible reactions, the lower bound –or in specific cases, the upper bound– was fixed at 0 mmol·gDW^−1^·h^−1^. Maximization of growth rate was set as the objective function. Anaerobic conditions were imposed by constraining the lower bound of the O_2_ exchange reaction to 0 mmol·gDW^−1^·h^−1^; whereas aerobic conditions were simulated by setting this bound to -20 mmol·gDW^−1^·h^−1^ (Nogales, 2017; Blázquez *et al*., 2023).

To prevent unrealistic behaviors under some conditions, certain reactions were deactivated and constrained to 0 mmol·gDW^−1^·h^−1^ as stated in reference *E. coli* models. Under all conditions, the following reactions were blocked: CAT, DHPTDNR, DHPTDNRN, SPODM, SPODMpp, SUCASPtpp, SUCFUMtpp, SUCMALtpp, and SUCTARTtpp, as well as PTRCORNt7pp, GALCTNLt2pp, GALCTLO, EX_chitob_e, URCN, CHITPH and CHITOBpts. The following reactions were kept active only in anaerobiosis: DTARTD, LCARS, SUCTARTtpp, TARTD, TARTRt7pp, ARGAGMt7pp, FHL and CELBpts (Monk *et al*., 2016). Although FHL is active anaerobically, it was also blocked here to allow quantification of formate production that would otherwise be fully consumed by the return reaction.

NGAM and GAM energy requirements were constrained according to the values specified in the previous *ATP expenditure parameters* section. For the simulation of each growth media, the lower bounds of exchange reactions were set to zero except for the metabolites present in the defined medium (M9 minimal media or gut microbiota media, GMM). Detailed specific exchange restrictions of each media are provided in Supplementary Methods. M9 minimal media was employed when simulating the experimental datasets, subject to additional condition-specific adjustments. A detailed summary of the constrained reactions and fluxes applied to in *silico* simulations of these datasets is provided in Supplementary Methods.

### Flux Balance and Flux Variability Analyses

Flux Balance Analysis (FBA) and Flux Variability Analysis (FVA) were performed using the COBRApy software (Ebrahim *et al*., 2013) All optimization problems were solved with CPLEX and the GNU Linear Programming Kit (GLPK). FBA was used to compute the maximum value of the objective function and the corresponding flux distribution (Thiele and Palsson, 2010; Orth *et al*., 2011). Maximization of the growth rate was set as the default cellular objective function. For modelling and simulating growth specific conditions, bounds of some reactions were modified as previously specified in *in silico constraints and growth conditions* section. Subsequently, FVA was carried out to determine the minimum and maximum feasible fluxes for each reaction on the constrained metabolic model while maintaining at least 90% of the optimal FBA objective value.

### Possibilistic Metabolic Flux and Consistency Analysis

Possibilistic Metabolic Flux Analysis (MFA) was applied as previously described in (Tortajada *et al*., 2010; Llaneras, 2011; Morales *et al*., 2014), using the PFAToolbox implemented in MATLAB R2020b. Briefly, this methodology accounts for data uncertainty, allowing experimental measurements to deviate from actual flux values, and can be represented as: wm = vm + em, where em is a vector containing the measurement errors. Each candidate solution fulfilling the model and experimental constraints is denoted as δ. A possibility value [0, 1] is assigned to each solution based on a linear cost index 𝐉(δ)=α·𝜀1+𝛽·𝜇1, where the possibility of each solution will be π(δ)=exp(−𝐉(δ)𝛿⊂Δ. Thus, a situation where vm = w is considered fully possible, and the greater the deviation between vm and wm, the lower the associated possibility.

In order to assess consistency, the agreement between experimental datasets and the metabolic network was evaluated as described in (Tortajada *et al*., 2010; Llaneras, 2011). Briefly, this method assigns a possibility value π^mp^ = [0, 1] to consistency between the most possible flux distribution with the model constraints and the experimental data under the model constraints. A value of 1 indicates full consistency, whereas lower values indicate discrepancies between measurements and model. For each dataset, measurement uncertainty was defined so that fluxes were considered fully possible (π = 1) within 5% of their reported standard deviations, while deviations exceeding the larger of either 20% of the experimental mean value or two standard deviations from experimental standard deviations corresponded to a possibility of π ≤ 0.1.

### Flux-associated error estimation

To quantify the agreement between experimental measurements and model-predicted fluxes, the Relative Squared Error (RSE) was computed. For each condition and model, individual flux RSE values were calculated and then summarized to obtain the median error estimate. These values were used to compare the predictive performance of the different EcN GEMs across experimental datasets.

### Metabolomic assays and data analysis

Starting from a 10 mL overnight culture in LB medium for each strain, the cells were washed twice with saline solution and inoculated into the corresponding medium at OD 0.05. Cultures were incubated overnight at 37°C under anaerobic conditions (BD GasPak™ system). After incubation, the cultures were collected and centrifuged for 15 minutes at 4500*g*. The supernatants were filtered through a 0.22 µm membrane filter and stored at -80°C until HPLC analysis. For each growth medium (GMM and M9), three replicates of both strains were performed, across two independent experimental setups. Blanks were conducted as medium samples, non-inoculated, independently.

The determinations of the metabolites listed in Supplementary Methods were carried out by high-performance liquid chromatography according to the analytical methods described below. For short-chain fatty acids (SCFAs) and fermentation-related metabolites, an Arc-HPLC system coupled with a refractive index detector (RID 2414) (Waters Corp., Milford, MA, USA) was used. The separation was carried out on an ion-moderated partition chromatography column (Aminex HPX-87H, 300 x 7.8mm, 9 µm) from Bio-Rad Laboratories (Hercules, CA, USA), kept at 60°C, and eluted with 5mM H_2_SO_4_, at 0,6 ml min^−1^ flow rate (Huda-Faujan, *et al*., 2010).

Amino acid quantification (arginine, ornithine, and glutamate) was carried out on a Waters Acquity Arc HPLC system equipped with a fluorescence detector (FLD 2475) using a Nova-Pak C18 column (3.9 × 150 mm, 4 µm). Analyses were performed at 37 °C with a 1 mL min⁻¹ flow rate following an AccQ-Tag gradient program. Detection was achieved at excitation and emission wavelengths of 250 nm and 395 nm, respectively. Standards and samples were derivatized according to the manufacturer’s instructions. Peak identification and quantification were carried out using analytical standards and their corresponding calibration curves (Jaworska *et al.,* 2012). Tryptophan and γ-aminobutyric acid (GABA) were quantified by UHPLC coupled to a single quadrupole mass detector (LC-QDa-MS) using a Waters Acquity Arc system with a Kinetex C18 column (100 × 4.6 mm, 2.6 µm) (Phenomenex, Torrance, CA, USA). Tryptophan was analyzed using water and methanol mobile phases containing 0.1% formic acid under a gradient elution at 0.5 mL min⁻¹, while GABA was measured under similar conditions with ammonium formate-formic acid buffers at 0.6 mL min⁻¹. Compounds were detected in positive electrospray mode in Single Ion Recording, and quantified using external calibration standards (Zhu, *et al*., 2011, de Bie, *et al*., 2021).

For data analysis and comparison, blanks measurements were extracted for each metabolite value, depending on medium. Metabolite concentrations (in mg/L) were normalized to cell growth in terms of OD. This processed data was used in the subsequent analysis. For visualization of each metabolite profiling, ggplot2 (v3.4.4; Wickham, 2009) was employed. Statistical analysis was performed in python (v3.11.5) using a bootstrap resampling with 10000 iterations and a significance level of 0.05 to evaluate differences between control and condition samples. A threshold of 10^−3^ mg/L was applied to filter out low-abundance metabolites. For each metabolite and condition, 95% bootstrap-based confidence intervals were computed to assess the significance of differences in metabolite concentrations relative to control values.

### Code availability

The code and data for reproducing this analysis are available in the GitHub repository: https://github.com/paocorbin/EcN-GEM

## 3. Results

### 3.1 Reconstruction of the EcN and structural comparison with other EcN GEMs

#### 3.1.1 Previously published GEMs of EcN

Several researchers have developed genome-scale metabolic models (GEMs) specific to EcN. The initial step in this research involved comparing the available reconstructions (Table 1) to evaluate how their structural features influence predictive accuracy and performance in practical experimental scenarios.

**Table 1.** Main features of published genome-scale metabolic models (GEMs) of *Escherichia coli* Nissle 1917. For each model, the table reports the total number of reactions, metabolites and genes, along with a brief description of its focus and the type of experimental validation used.

| Model ID | CANYUN model | iDK1463 | iHM1533 |
| --- | --- | --- | --- |
| Reference | Moutinho *et al.*, 2022 | Kim *et al.*, 2021 | van't Hoff *et al.*, 2022 |
| Reactions | 737 | 2984 | 3143 |
| Metabolites* | 2607 (1661) | 2112 (1313) | 2261 (1440) |
| Genes | 277 | 1464 | 1533 |
| Experimental validation | Phenome analysis in aerobic and anaerobic conditions with Biolog assays for carbon sources. | Phenome analysis in aerobic conditions with Biolog assays for carbon, nitrogen, phosphorus, sulfur, nutrient supplements and inhibitory substrates. | Phenome analysis in aerobic conditions with Biolog assays for carbon, nitrogen, phosphorus and sulfur sources. Previously published <sup>13</sup>C fluxomics data. |
| Focus | CANYUN method validation and example of utility. | Comparative metabolic traits with other *E. coli* strains relevant to commensalism and gut colonization. | Optimization of secondary metabolites production. |

\*Numbers in parentheses indicate non-redundant metabolites, counted only once regardless of their compartmental location.

The first GEM of EcN (iDK1463; Kim *et al*., 2021) was derived from earlier reconstructions of *E. coli* K-12 MG1655 and two uropathogenic strains, CFT073 and ABU83972. This model was qualitatively validated using Biolog phenotypic assays, covering a broad range of carbon and nitrogen sources, and other substrates under aerobic conditions. The analysis identified gene clusters unique to EcN relative to the K-12 strain and used the metabolic model to simulate anaerobic growth in intestinal environments with various carbon sources. These simulations revealed distinct metabolic behaviors between EcN and K-12 strains.

Subsequently, Moutinho and coworkers (2022) reconstructed a smaller, more streamlined network to demonstrate their proposed reconstruction workflow, called Constraint-based Analysis

Yielding reaction Usage across metabolic Networks (CANYUN). This model was validated against experimental growth data under both aerobic and anaerobic conditions.

Finally, van’t Hof and coworkers (2022) developed an additional model (iHM1533), built on a high-quality genomic sequence and related to 55 other *E. coli* strains with available GEMs. This model was validated through phenome analysis and ^13^C-fluxomics data, which provided insights into internal fluxes under aerobic conditions (Revelles *et al*., 2013). They also refined the pathways involved in secondary metabolism to create a framework for using EcN as a synthetic biology chassis, particularly for enhancing secondary metabolite production, such as antimicrobial compounds. For this, a mutant EcN strain curated from the cryptic plasmids was used for further validation during Biolog phenotypic assays.

An analysis of the main features of these EcN models shows an average of approximately 3,064 reactions (excluding the Canyun model) and around 1,471 unique metabolites. The Canyun model differs from the others because its primary goal was to simulate qualitative growth phenotypes and validate the reconstruction workflow. As a result, it omits over 1,200 reactions that are not essential for growth. Overall, the published EcN GEMs vary in complexity and the level of manual curation, which depends on the reconstruction strategy—whether based on extensive reference GEMs or utilizing new reconstruction methods—and the degree of refinement performed. The models listed in Table 1 are ranked from least to most complex based on these criteria. Nevertheless, all reconstructions have been validated using similar experimental datasets.

#### 3.1.2 Development of the EcN model presented in this study

In this study, we present an updated GEM of EcN, designated iPC1373 (Supplementary Material S1; Alvarez *et al*., 2019). The model was developed to accurately represent the core physiology and metabolic capabilities of this probiotic strain. Reconstruction was based on the annotated genome of EcN, available GEMs of closely related *E. coli* strains, and manual curation.

Initially, a draft reconstruction was generated by incorporating all shared reactions and metabolites from the *E. coli* K-12 MG1655 and the uropathogenic *E. coli* CFT073 models, based on the presence of homologous genes. Manual curation was subsequently performed to refine the model, addressing non-homologous genes, filling metabolic gaps, verifying reaction directionality, and removing redundant reactions based on literature evidence. The final step involved the annotation of reactions, metabolites, and genes using multiple databases, including KEGG, BioCyc, MetaNetX, SEED, Rhea, ChEBI, and NCBI. Systems Biology Ontology (SBO) terms were assigned using SBOannotator (Leonidou *et al*., 2023).

Unlike previously published models, which do not include the EcN cryptic plasmids pMUT1 and pMUT2, iPC1373 explicitly incorporates these genetic elements. The concept of plasmid-associated metabolic burden is well established, and several studies have addressed its representation in metabolic models of various scales, including GEMs (Ow *et al*., 2009; Zeng and Yang, 2019; Oftadeh and Hatzimanikatis, 2024). Notably, simulations of wild-type EcN growth on glucose using previously published models overestimated the experimental biomass growth rate by approximately 20% (van’t Hof *et al*., 2022), suggesting an unaccounted burden or metabolic cost. To capture this effect, iPC1373 includes the maintenance and expression of both cryptic plasmids, implemented as two additional reactions (Eq. 1 and Eq. 2) representing their associated biosynthetic and energetic demands. Plasmid maintenance costs were parameterized using literature-derived estimates for plasmid-bearing *E. coli* (Weber *et al*., 2002; Orth *et al*., 2011; Zeng and Yang, 2019). Accordingly, the NGAM parameter was adjusted by constraining the lower bound of the ATPM reaction to the median of reported values: 25 mmol·gDW^−1^·h^−1^ under aerobic conditions and 3.5 mmol·gDW^−1^·h^−1^ under anaerobic conditions. For additional validation, the effect of ATP parameters on reaction fluxes, particularly biomass estimation, was evaluated (see Supplementary Methods). A good correspondence was found between the values selected from literature, and the resulting predictions using biomass as the objective function.

The finalized model comprises 2,716 reactions, 1,943 metabolites (corresponding to 1,217 unique metabolites), and 1,373 genes. Model quality and consistency evaluated using the Memote testing suite (Lieven *et al*., 2020), yielded a total score of 91% and a consistency score of 98% (Supplementary Material 2). Annotation completeness scores were 78% for metabolites, 79% for reactions, and 60% for genes; the latter reflecting incomplete database coverage. The coverage of SBO terms reached 93%, indicating a high level of semantic and structural annotation.

#### 3.1.3 Structural analysis and comparison of available EcN metabolic models

Once the iPC1373 model had been established, we evaluated its structural correspondence with previously published reconstructions. The EcN CANYUN model was excluded from the comparison and subsequent analyses owing to its deliberately reduced size. In terms of reactions (Fig. 1A, Supplementary Material 3), the models shared 2,654 reactions, 83% of the total. Notably, the published models iDK1463 and iHM1533 contain substantially more reactions; over half of their surplus (154 of 284; 58%) comprises extracellular exchange and transport reactions, followed by 39 reactions involved in alternative carbon metabolism (Fig. S1). The iPC1373 model exhibits a smaller set of 17 unique reactions, comprising: i) two biomass formulations—wt and core, corresponding respectively to the full, experimentally measured macromolecular composition of the wild-type strain and a simplified precursor-based biomass reaction—, ii) the two reactions related to plasmid representation, iii) seven transport reactions, and iv) six reactions representing the O-antigen biosynthesis module (Wzx/Wzy/WaaL pathway), responsible for semirough LPS production in EcN (Grozdanov *et al*., 2002). In iDK1463, apart from the biomass reaction, the other 19 unique reactions were removed during the curation of iPC1373 as duplicates or redundancies—changes consistent with the updated *E. coli* K-12 model iML1515 (e.g., ALAt2pp, ALAt2pp_copy1, ALAt2pp_copy2). Finally, iHM1533 includes 188 unique reactions (Fig. S2), primarily in secondary-metabolite biosynthesis (102), transport and exchange (43), and aromatic compounds breakdown (15).

**Figure 1.**
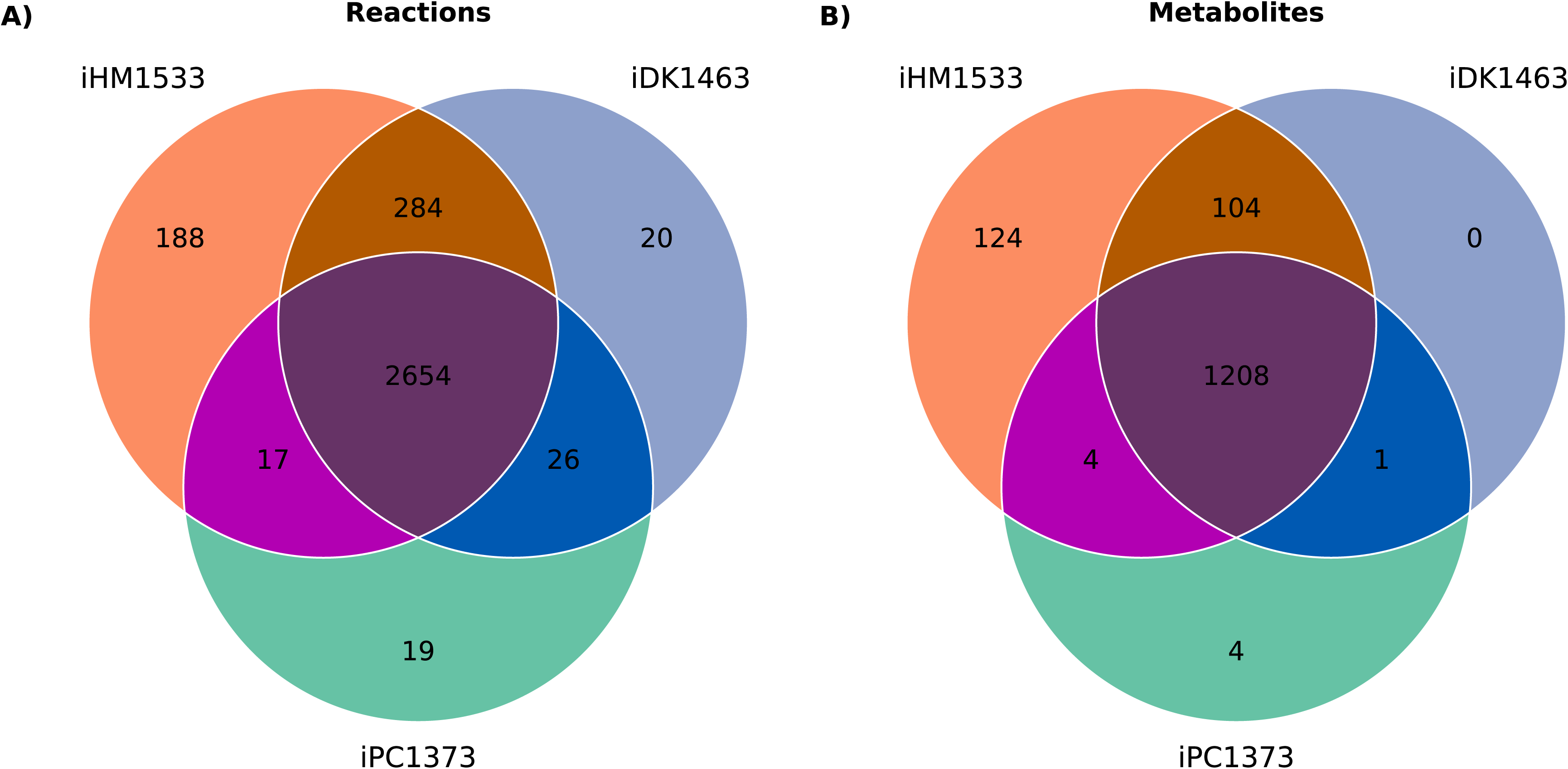
Comparison of reactions (A) and metabolites (B) among available *Escherichia coli* Nissle 1917 GEMs.

For metabolites (Fig. 1B, Supplementary Material 4), the main differences occur in iHM1533 and reflect the subsystem expansions aimed at optimizing specialized metabolism (Fig. S3), consistent with the reaction-level disparities among the models. In terms of MEMOTE (Fig. S4), the three EcN models show high consistency (97–98%). Annotation coverage is strong for metabolites and reactions (78–84%), with gene annotation being the main weakness (60–68%). iPC1373 has the highest SBO term coverage (93%). Overall scores are high and comparable (89–91%), with iPC1373 slightly leading by two points, indicating all three reconstructions are robust with minor annotation differences.

### 3.2 Model validation with experimental flux datasets

The next step was to validate the phenotypic consistency of the EcN metabolic models against independent experimental data, beyond their mathematical accuracy as calculated with MEMOTE. Equations reflecting plasmid replication were incorporated into all EcN models available for validation, comparison and predictive analyses. Unless otherwise specified, models including these plasmid-associated reactions were employed.

A quantitative approach was adopted to rigorously evaluate model performance of the different EcN metabolic models, examining whether increased model complexity improves quantitative predictive accuracy (O’Brien *et al*., 2015; Blázquez *et al*., 2023). Since completeness and mathematical consistency do not necessarily ensure accurate physiological predictions, this validation aimed to test the models’ ability to reproduce experimental biomass formation, substrate uptake, and by-product secretion rates—key parameters for bioprocess control and understanding EcN physiology. For that, several datasets from the literature were compiled (see above *Experimental datasets* in Methods).

#### 3.2.1 Experimental growth scenarios and model consistency analysis

First, the experimental datasets were used to evaluate consistency—that is, the compatibility between the experimental data and the model—by applying possibilistic MFA and computing the possibility (π) of the most possible flux state. As shown in Table 2, the consistency analysis yielded possibility values of 1 for the Revelles and coworkers (2013) dataset, indicating full agreement between the data and the models. For both fucose and rhamnose conditions, consistency was marginally higher in the iHM1533 model, increasing with model complexity. Conversely, randomly generated scenarios yielded possibilities lower than 0.1 on over 98% of occasions.

**Table 2.** Experimental data for EcN cultures and model consistency. Consistency values (π) were calculated using the possibilistic MFA method for each dataset across the evaluated models: iPC1373, iDK1463, and iHM1533. The data shown are the specific growth rate (h^−1)^, substrate uptake (q_S_), acetate formation (q_ace)_, pyruvate formation (q_pyr_), CO_2_ formation (q_CO2_), glycogen formation (q_glyc_) and 1,2-propanediol formation (q_12ppd_) obtained from datasets *(Revelles *et al*., 2013) and **(Millard *et al*., 2021) under aerobic conditions.

| Condition | $\mu$ | $q_s$ | $q_{ace}$ | $q_{CO_2}$ | $q_{pyr}$ | $q_{glyc}$ | $q_{12ppd}$ | Consistency | | |
| --- | --- | --- | --- | --- | --- | --- | --- | --- | --- | --- |
| | (h<sup>-1</sup>) | mmol·gDW<sup>-1</sup>·h<sup>-1</sup> | | | | | | $\pi$<br>iPC1373 | $\pi$<br>iDK1463 | $\pi$<br>iHM1533 |
| Glucose wt* | 0.79<br>±0.02 | 12.50<br>±1.00 | 6.30<br>±0.10 | — | n.d | 0.003<br>±0.010 | — | 1 | 1 | 1 |
| Gluconate wt* | 0.74<br>±0.02 | 20.00<br>±1.70 | 10.40<br>±0.10 | — | 3.20<br>±0.10 | 0.001<br>±0.010 | — | 1 | 1 | 1 |
| Glucose $\Delta csrA51$* | 0.63<br>±0.20 | 10.80<br>±1.00 | 7.20<br>±0.10 | — | n.d. | 0.057<br>±0.014 | — | 1 | 1 | 1 |
| Gluconate $\Delta csrA51$* | 0.57<br>±0.02 | 12.60<br>±1.70 | 6.30<br>±0.10 | — | 0.40<br>±0.10 | 0.007<br>±0.010 | — | 1 | 1 | 1 |
| Fucose** | 0.48<br>±0.01 | 12.80<br>±0.30 | 4.00<br>±0.20 | 15.30±<br>2.40 | — | — | 11.10±<br>0.20 | 0.32 | 0.35 | 0.45 |
| Rhamnose** | 0.28<br>±0.01 | 6.90<br>±0.20 | 0.20<br>±0.10 | 12.30±<br>0.70 | — | — | 5.50<br>±0.10 | 0.41 | 0.45 | 0.60 |
| Random glucose | 0-10 | 0-10 | 0-10 | — | 0-10 | 0-10 | — | < 0.1:<br>98% | < 0.1:<br>99% | < 0.1:<br>99% |

*Random datasets* correspond to 100 automatically generated flux datasets used for consistency testing.
n.d., not detected; —, not measured.

#### 3.2.2. Prediction of biomass growth rate

Then, we evaluated whether the incorporation of the plasmid module and the adjustment of energy maintenance rates improved the prediction of biomass growth rates across the experimental datasets. Introducing the plasmid module partially corrected the overestimation of *in vivo* growth observed in all simulated carbon sources (Table 3). Model predictions including the plasmid representation showed better agreement with experimental data, clearly decreasing the relative squared error (RSE) across all scenarios and models, resulting in an average reduction from 16% to 4% (Table S1). There were no wide differences in performance between the different models, with the iDK1463-based reconstruction achieving the best biomass prediction, with a prediction error of 3.6%, compared to 4.0% and 4.6% for iHM1533 and iPC1373, respectively.

**Table 3.** Comparison between experimental and model-predicted biomass growth rates. For each experimental dataset, all measured external fluxes—except biomass—were constrained to their experimental values within the standard deviation. The table shows the FBA-optimized growth rate predicted for each model, evaluated both with and without plasmid consideration.

| Models excluding plasmid consideration |  |  |  |  |
| --- | --- | --- | --- | --- |
|  | Exp. | iPC1373 | iDK1463 | iHM1533 |
| Condition | Growth rate (h<sup>-1</sup>) |  |  |  |
| Glucose wt | 0.790 | 1.098 | 1.059 | 1.097 |
| Gluconate wt | 0.740 | 1.103 | 1.064 | 1.105 |
| Glucose ΔcsrA51 | 0.630 | 0.893 | 0.881 | 0.914 |
| Gluconate ΔcsrA51 | 0.570 | 0.958 | 0.925 | 0.961 |
| Fucose | 0.480 | 0.493 | 0.493 | 0.513 |
| Rhamnose | 0.280 | 0.334 | 0.335 | 0.348 |
| Models including plasmid consideration |  |  |  |  |
|  | Exp. | iPC1373 | iDK1463 | iHM1533 |
| Condition | Growth rate (h<sup>-1</sup>) |  |  |  |
| Glucose wt | 0.790 | 0.903 | 0.870 | 0.902 |
| Gluconate wt | 0.740 | 0.914 | 0.882 | 0.916 |
| Glucose ΔcsrA51 | 0.630 | 0.816 | 0.805 | 0.836 |
| Gluconate ΔcsrA51 | 0.570 | 0.771 | 0.743 | 0.771 |
| Fucose | 0.480 | 0.388 | 0.390 | 0.406 |
| Rhamnose | 0.280 | 0.236 | 0.238 | 0.248 |

#### 3.2.3 Prediction of remaining external rates

Once plasmid incorporation was shown to improve biomass growth prediction, external production rates were estimated using the plasmid-bearing models by constraining only the carbon source uptake rate to the experimental values of each dataset. This approach reflects typical fermentation conditions, where substrate uptake is measured and controlled, while other exchange fluxes are rarely monitored. Predictions for all datasets were obtained using FBA, FVA, and possibilistic MFA (Figure 2; Supplementary Material S5).

**Figure 2.**
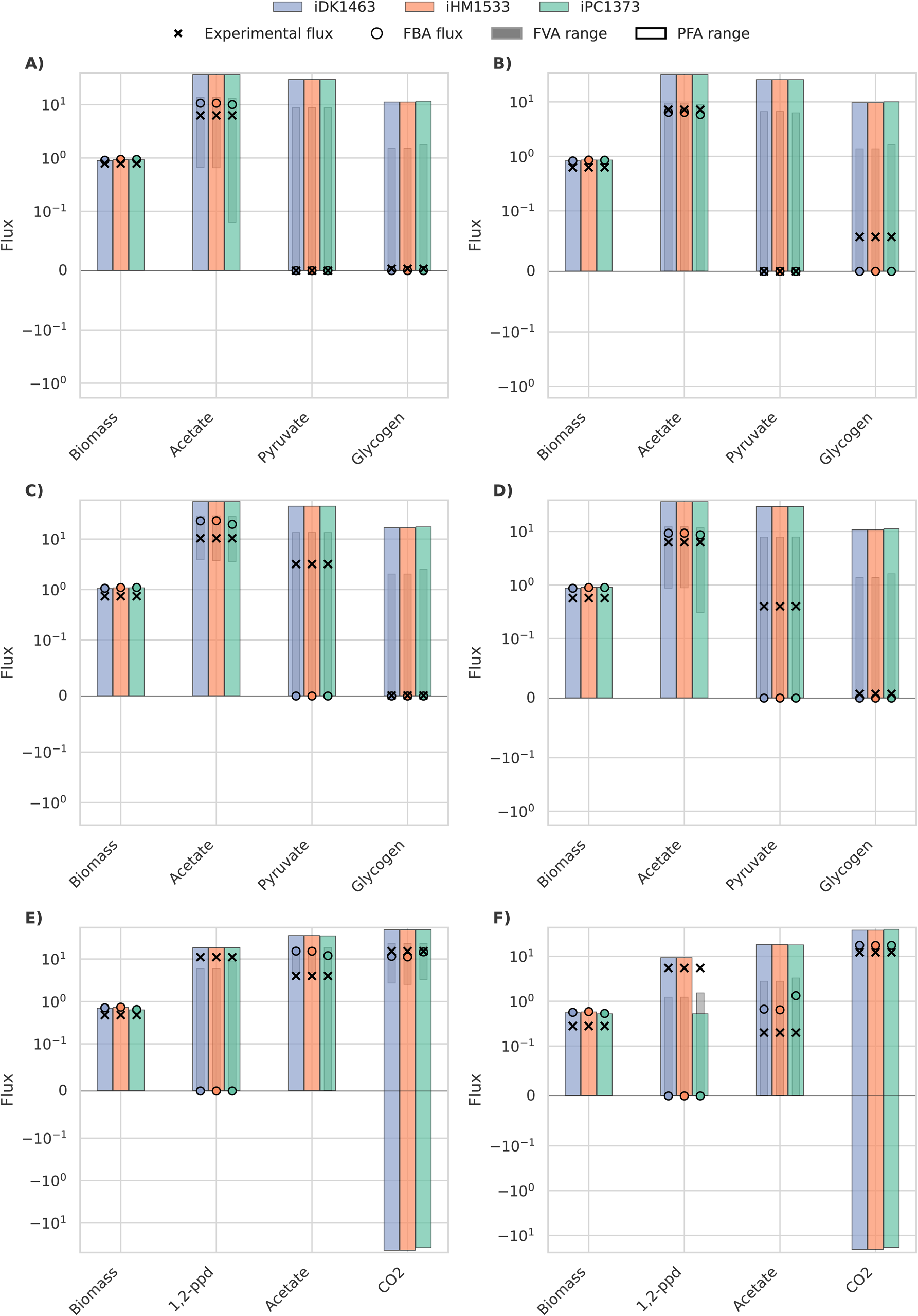
External fluxes predicted by the models iDK1463 (blue), iHM1533 (orange), and iPC1373 (green). Simulations were performed under aerobic conditions by constraining substrate uptake rates to the corresponding experimental values. Panels A and B show predictions for glucose in the wild type and the ΔcsrA51 mutant strains, respectively; panels C and D show predictions for gluconate in the wild type and the ΔcsrA51 mutant; panel E and F corresponds to fucose and rhamnose as sole carbon sources, respectively. Gray internal bars indicate FVA ranges; colored bars (in blue, green and orange) represent the range of possibilistic MFA solutions; circles denote FBA solution; and crosses mark experimental flux measurements. Fluxes are displayed using a symlog scale with a linear threshold of 0.01 to highlight differences at low magnitudes. Original flux values are provided in Supplementary Material S5.

Overall, a good directional agreement was observed between experimental and *in silico* FBA-predicted values (Table S2), though quantitative deviations were larger than those found for biomass estimation, with an overall average RSE of 32% (Table S3). Among the models, iPC1373 provided in this case the most accurate predictions, achieving an RSE of 27%, compared to 35% for both iDK1463 and iHM1533.

Additionally, the three models produced comparable external flux predictions across all methodologies, with only minor variations. Noticeable discrepancies appeared for 1,2-propanediol production (Fig. 2E, Fig. 2F), pyruvate production under gluconate conditions (Fig. 2C, Fig. 2D), and glycogen formation (Fig. 2B). Range-based approaches (possibilistic MFA and FVA) generally encompass both the FBA predictions and the experimental values within their bounds. In some cases—such as 1,2-propanediol production—these range-based methods captured the experimental results more accurately.

#### 3.2.4 Prediction of internal flux reactions rates

Finally, since the complete flux distribution can be estimated, we compared the model predictions with the experimental ^13^C fluxomics data reported for the EcN wild-type strain by Revelles and coworkers (2013), using glucose (Fig. 3A) and gluconate (Fig. 3B) as sole carbon sources. Both FBA and possibilistic MFA were applied to analyze central metabolic pathways and reactions (Fig. 3, Supplementary Material S6), allowing for a comparison of the methodologies. Overall, model predictions qualitatively followed the experimental intracellular flux trends, except for the TCA cycle, which showed underestimated fluxes under both carbon sources–except for the iHM1533 model in glucose using possibilistic MFA–and reaction 7 of glycolysis (the conversion of phosphoenolpyruvate to pyruvate) under glucose. Quantitative discrepancies depended on both the model and the methodology, although FBA predictions were generally accurate across reaction groups. However, notable deviations were observed in reactions 4 (glucose) and 14 (gluconate), where the iPC1373 and iHM1533 models demonstrated greater consistency with the experimental findings. Overall, attending to median RSE, all models performed similarly across the different scenarios with relative errors ranging 29-33% (Table S4).

**Figure 3.**
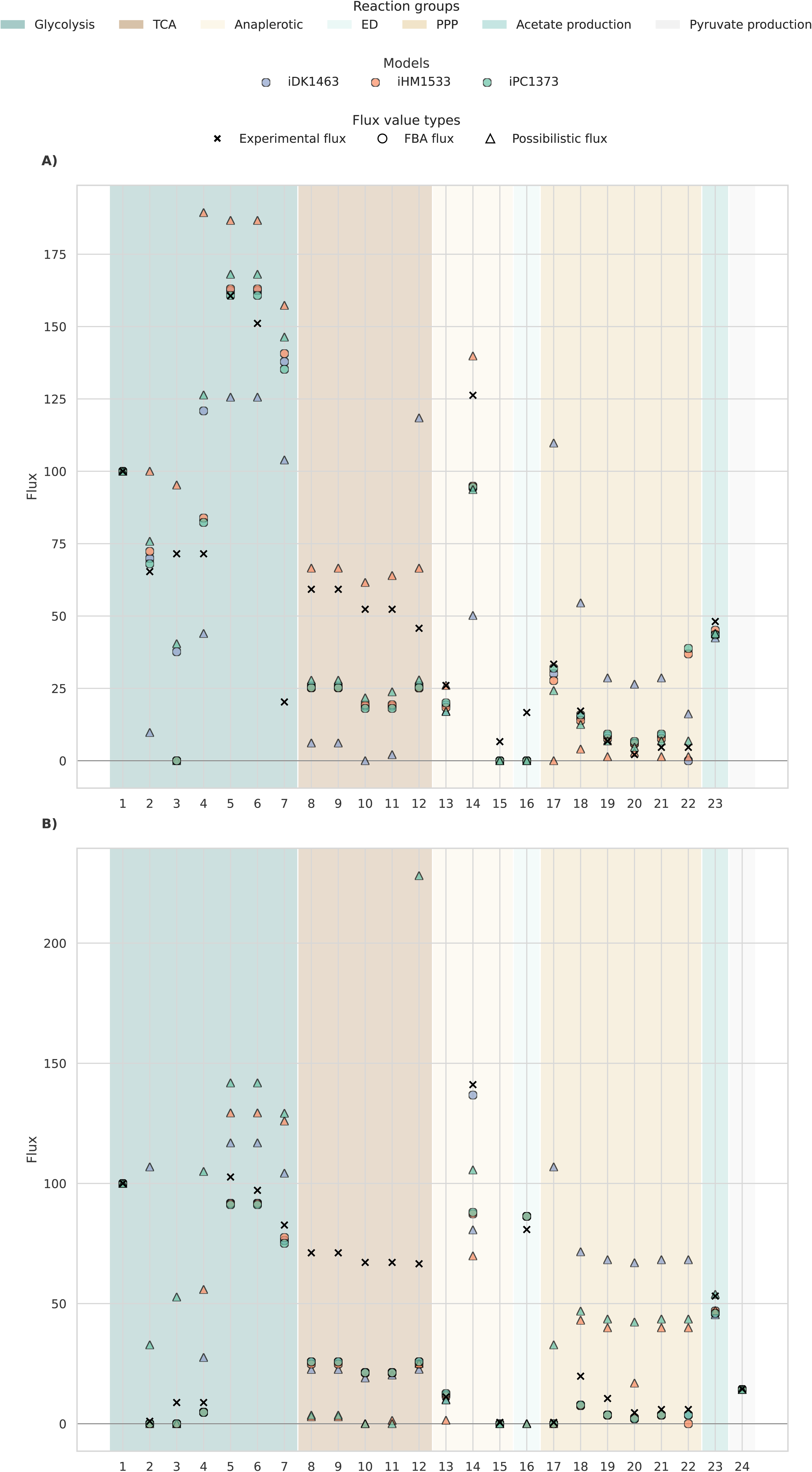
Predicted flux distributions in central metabolism for the models iDK1463 (blue), iHM1533 (orange), and iPC1373 (green) under wildtype conditions using glucose (A) and gluconate (B). Simulations were performed by constraining the measured exchange fluxes to their experimental values (Revelles et al., 2013). Circles denote FBA solution; triangles represent possibilistic MFA at *π* = 1; and black crosses indicate experimental ^13^C flux measurements. Fluxes are shown on a linear scale and normalized to the carbon source uptake rate (set to 100). Flux values can be found in Supplementary Material S6. Central metabolism reactions are grouped as: glycolysis/gluconeogenesis (Glycolysis), tricarboxylic acid (TCA) cycle, anaplerotic reactions, Entner-Doudoroff (ED) pathway, pentose phosphate pathway (PPP), and acetate/pyruvate production. Reaction indices (1–24) correspond to the entries in Table S5, which list the metabolite conversions (Revelles *et al.,* 2013) and their corresponding BiGG database reaction IDs.

When possibilistic MFA was applied to estimate the most probable flux value for each reaction, larger differences emerged depending on the underlying model. In this case, the model iPC1373 achieved the highest overall accuracy, better capturing the fluxes through the TCA cycle reactions and exhibiting the lowest median RSE across the 23-24 reactions analyzed, 32% (Table S4).

### 3.3 EcN exhibits a distinct metabolomic profile in GMM medium, both experimentally and *in silico*

Once the models were validated and compared against experimental flux datasets, we examined whether they could reproduce EcN’s metabolic behavior under different environmental conditions. To this end, EcN and the reference *E. coli* K-12 strain were cultured anaerobically in minimal M9 media and Gut Microbiota Medium (GMM), which promotes the growth of beneficial gut bacteria and enhance short-chain fatty acids (SCFAs) production (Park *et al*., 2024). This setup aimed to determine if GMM elicits distinct metabolic behavior in the probiotic strain, as it mimics the nutrient composition of the gut environment. Although previous GEM studies (Kim *et al.,* 2021) suggested that EcN behaves differently under anaerobic environments, this physiologically relevant aspect has not yet been experimentally confirmed best of our knowledge.

To explore global metabolic differences, representative metabolites–including organic acids, SCFAs, and amino acids–were quantified. Samples were grouped by strain and medium, revealing largely similar profiles overall, except for EcN grown in GMM, which clearly separated from the rest (Fig. 4A). This separation was mainly driven by PC1, which explained 82% of the total variance, followed by PC2 (10.5%). The metabolites contributing most to this separation were lactate, acetate, formate, and succinate.

**Figure 4.**
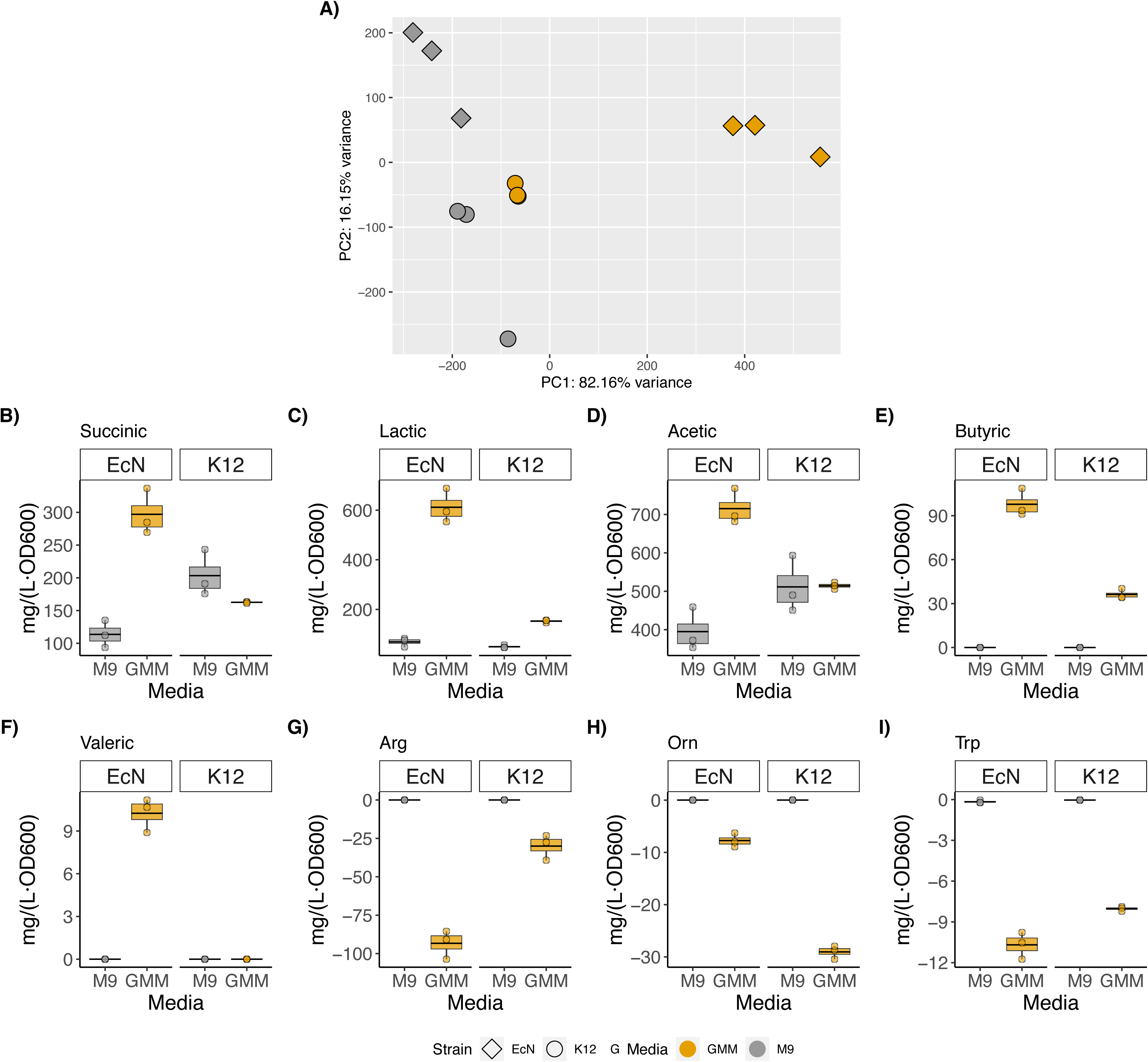
Metabolite profiling of *E. coli* Nissle (EcN) and K-12 in M9 medium with glucose and Gut Microbiota Medium (GMM). A) Principal Component Analysis of HPLC quantified metabolites for *E. coli* K-12 (circle) and EcN (diamond) cultured in GMM (yellow) or M9 (gray) medium. B-I) Boxplots showing metabolite levels (in mg/L·OD600) grouped by strain and medium. Each group includes three biological replicates, displayed as individual points (circles) within the boxplots. Succinic, lactic, acetic, butyric and valeric acids correspond to SCFAs and fermentation-related metabolites; while Arg (arginine), Orn (ornithine), Trp (tryptophan) are amino acids. Positive values correspond to production of metabolites and negative, to consumption.

When analyzing specific patterns in the different extracellular compounds, some metabolites showed no detectable production or consumption or did so only in insignificant changes in several compounds, including propionate, isobutyrate, and isovalerate (Supplementary Material S7). In contrast, clear differences were observed in the production of organic acids—particularly SCFAs—and in amino acid consumption when EcN was grown in GMM medium (Fig. 4B-I). Indeed, an overall increase in fermentation-related metabolites and SCFAs was found in EcN cultures under GMM conditions (Fig. 4B-F). EcN produced acetic, butyric, formic, lactic, and succinic acids at net concentrations of 1.19, 0.16, 0.49, 1.01, and 0.49 g/L, respectively (Supplementary Material S7). In addition, small amounts of valeric acid (≈20 mg/L) were detected exclusively in EcN cultures (Fig. 4F). Compared to the K-12 strain, EcN showed markedly higher production of acetic, butyric, lactic, and succinic acids (0.08, 0.08, 0.33, and 0.35 g/L in K-12, respectively). Moreover, EcN demonstrated distinct amino acid utilization patterns within GMM (Fig. 4G-I), exhibiting elevated uptake of arginine and tryptophan compared to the K-12 strain or EcN in M9 medium. Interestingly, ornithine consumption showed opposite behavior for the strains. These observed variations seem independent of growth, as EcN achieved a lower optical density under these conditions (Supplementary Material S7).

Finally, the models were used to simulate EcN metabolism under M9 and GMM media (Fig. 5, Supplementary Material S8). As previously evaluated, all three models successfully captured the overflow metabolism characteristic of EcN growth in M9 medium with glucose (Fig. 5A), including the production of acetate, ethanol and formate, although they failed to reproduce lactate formation. Under simulated GMM conditions (Fig. 5B), all three models accurately reproduced the distinctive overproduction of succinate and acetate observed experimentally. Model iPC1373 also captured the increased butyrate production, while iHM1533 predicted higher lactate formation. In contrast, ethanol production was not predicted by any model—which aligns with its low experimental levels when compared to M9 medium (Supplementary Material S7). Enhanced consumption of key amino acids, including glutamate, arginine, and tryptophan, was also correctly represented. However, none of the models reproduced ornithine uptake, with iHM1533 instead predicting its secretion. Some discrepancies, such as the absence of isovaleric, valeric, and GABA predictions were probably due to missing reactions or metabolites in the reconstructions.

**Figure 5.**
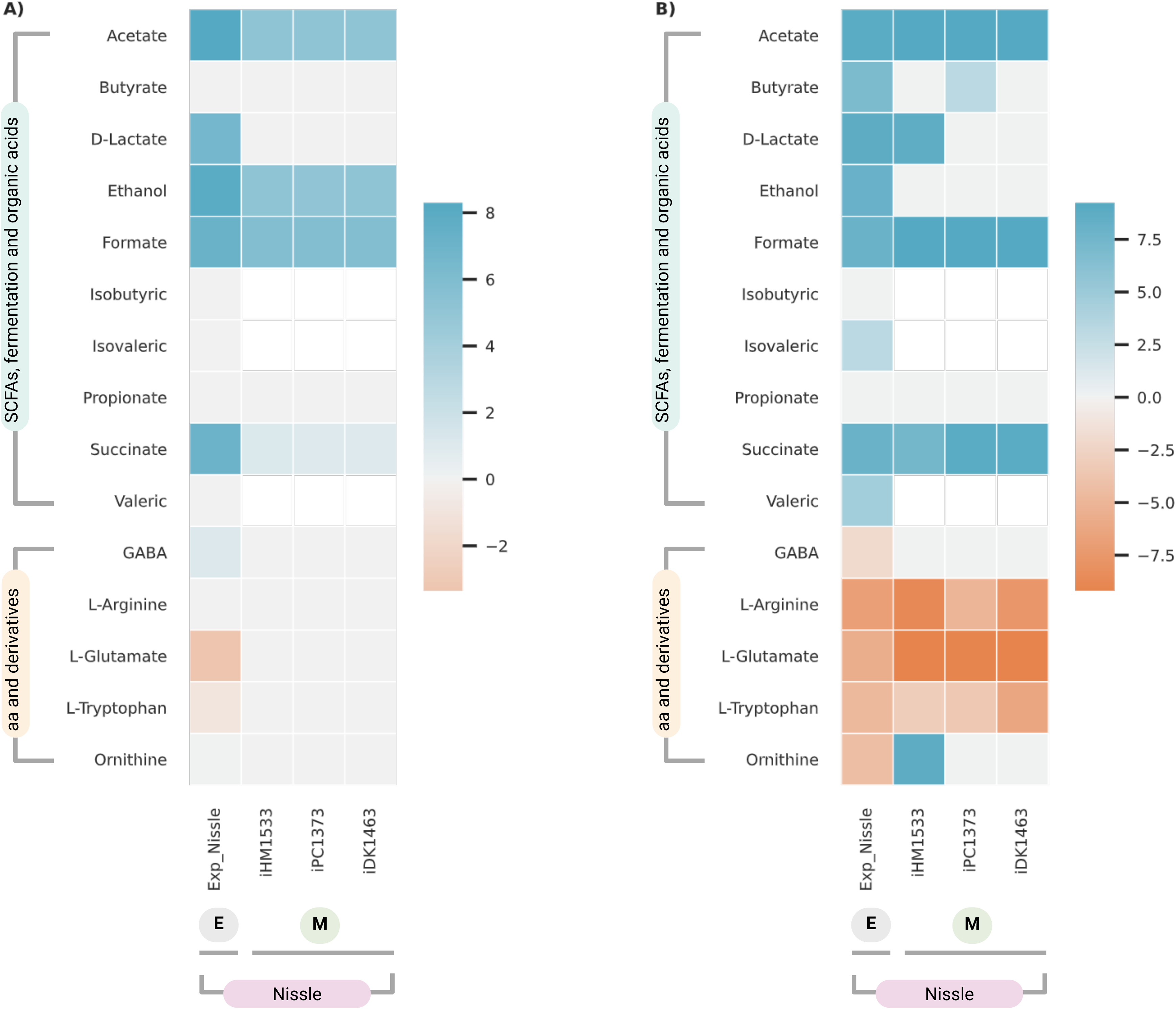
Simulation results of EcN models under M9 medium with glucose and gut microbiota medium (GMM). The heatmap shows the FBA predicted fluxes for models iHM1533, iDK1463 and iPC1373 under the *in silico* conditions of M9 medium with glucose (A) and GMM medium (B), along with the corresponding experimental results for the EcN strain. Data are displayed using a symlog scale to facilitate comparison between experimental measurements and model predictions, including only those metabolites for which both experimental and simulated values are available. The scale indicates consumed metabolites in orange, produced metabolites in blue, and those with a value of zero –neither consumed nor produced– in gray. Experimental values are provided in Supplementary Material S7; original flux values, in Supplementary Material S8.

## 4. Discussion

The main objective of this study was to develop a GEM of EcN capable of accurately reproducing its physiological behavior, with particular focus on its characteristic probiotic traits. Although EcN is a well-characterized and commercially available probiotic strain, systems biology studies have paradoxically emphasized its potential as a biosynthetic chassis rather than its probiotic physiology. Owing to its strong adherence, resistance, and persistence in the gastrointestinal tract, EcN represents an attractive platform for targeted drug delivery and live biotherapeutic applications (Kan *et al.,* 2021; Kim *et al.,* 2021). Previous research has typically relied on specific experimental constraints, media compositions, or genetically modified variants—such as the plasmid-cured strain reported by van’t Hof *et al*. (2022)—to characterize EcN’s performance as a recombinant host.

In constructing our model, we explicitly incorporated the presence and energetic cost of EcN’s cryptic plasmids. The metabolic burden associated with plasmid maintenance is well established in *E. coli* (Ow *et al*., 2009; Oftadeh and Hatzimanikatis, 2024), and previous EcN GEMs have tended to overestimate biomass growth rates (van’t Hof *et al*., 2022). Van’t Hof and colleagues attributed these discrepancies to incomplete biomass formulations, underestimated maintenance energy requirements, or missing by-product pathways. By introducing a plasmid-specific module representing the additional energetic demands of plasmid maintenance, our model achieved more realistic biomass yield estimates and accurately reproduced acetate overflow metabolism. This adjustment reduced the overestimation of *in silico* growth rates to within 5% of experimental measurements. Beyond improving model fidelity, this refinement is also relevant to synthetic biology applications, where EcN’s cryptic plasmids are often retained and exploited as stable endogenous expression vectors. The systematic inclusion of endogenous plasmids could also have broader implications for GEM development in other microorganisms. Many industrially and clinically important bacteria—whether probiotic, pathogenic, or commensal—naturally harbor plasmids (Fogarty *et al*., 2024; Garcillán-Barcia *et al.,* 2025), yet these elements are rarely represented in their respective GEMs. Incorporating plasmid-associated metabolic costs and interactions thus represents a promising strategy to enhance both model realism and predictive power.

To further assess model performance, we compared several EcN GEMs after integrating the plasmid module, evaluating the trade-offs between model complexity and accuracy in reproducing experimental physiological behavior. Interestingly, models with differing levels of curation and structural detail yielded distinct quantitative outcomes when validated against experimental flux data. Notably, greater model complexity did not necessarily improve predictive accuracy—particularly for external fluxes such as acetate or ethanol secretion—and similar inconsistencies were observed for intracellular flux predictions. Some recurring discrepancies, including those related to 1,2-propanediol, pyruvate, or glycogen production, likely reflect inherent limitations of FBA, which assumes growth optimization and neglects non-essential or suboptimal fluxes. These by-products are often regulated through mechanisms such as CsrA-dependent glycogen accumulation, underscoring the need to integrate regulatory constraints or multi-objective formulations to more faithfully reproduce EcN’s observed metabolism.

Our analyses further demonstrated that intracellular flux prediction accuracy does not scale linearly with model complexity. Predictive performance depended on both the computational method and the reaction–model pairing, indicating that specific models may better capture particular pathway dynamics. Under certain environmental simulations, some metabolites—such as valerate, isovalerate, and GABA—were not predicted by all models, likely due to missing exchange reactions or incomplete pathway representations. Other deviations, such as the absence of lactate production in IDK1463 and iPC1373 or misrepresented ornithine flux in iHM1533, may result from alternative flux distributions that divert carbon flow from these pathways.

While qualitative predictions aid in interpreting general physiological trends, quantitative precision is crucial for industrial applications. In probiotic manufacturing, accurate control of fermentation parameters is essential to maintain cell viability and product stability. GEMs are valuable tools for optimizing such processes, especially when it is difficult the direct measurement of key parameters such as growth, substrate uptake, or by-product secretion. Reliable model predictions can thus improve fermentation efficiency and product consistency. Accordingly, the present study establishes a framework for developing and validating strain-specific GEMs with quantitative data and industrial relevance.

From a methodological perspective, this work also illustrates how distinct metabolic flux analysis approaches provide complementary insights into cellular metabolism. Possibilistic MFA proved particularly valuable for incorporating measurement uncertainty and generating robust flux estimates in underdetermined systems, where conventional MFA or FBA may produce less stable solutions. Comparative analyses of FBA, FVA, and possibilistic MFA revealed that each method captures unique aspects of metabolic behavior and uncertainty. Furthermore, as shown in this study, different metabolic reconstructions—regardless of their origin or level of curation—can be effectively prioritized depending on the specific biological or experimental context.

Finally, this study enhances our understanding of EcN’s unique probiotic features. Metabolomic analyses confirmed that EcN displays a distinctive metabolic profile under anaerobic conditions in GMM, supporting previous *in silico* observations that it expresses unique metabolic traits under gut-like environments (Kim *et al*., 2021; van’t Hof *et al*., 2022). This phenotype is characterized by selective consumption and secretion of key metabolites that distinguish EcN from other *E. coli* strains. Compared to *E. coli* K-12, EcN consumes several amino acids at higher rates, potentially conferring a competitive advantage in the intestinal niche by depleting local nutrient pools and limiting resources available to competing microorganisms. Simultaneously, EcN produces elevated levels of organic acids—such as lactic and acetic acid—that inhibit pathogens, as well as SCFAs like butyrate and valerate, which promote the growth of beneficial anaerobes. Together, these findings reinforce EcN’s physiological distinctiveness and provide ecological insight into its probiotic mechanisms of action. They also confirm that EcN’s phenotype is strongly context-dependent, particularly on growth medium composition and oxygen availability. Thus, physiologically relevant conditions are essential when validating and applying GEMs to commensal, probiotic, or pathogenic microorganisms.

Future work could extend this framework to simulate microbial interactions within the gut ecosystem. Integrating EcN into community-scale metabolic models—similar to prior approaches for *Bacillus* and *Lactobacillus* species (Devika *et al*., 2021)—would allow exploration of its interactions with both commensal and pathogenic bacteria, offering deeper insights into its ecological role and therapeutic potential.

## 5. Conclusions

In summary, this study presents a refined genome-scale metabolic model of EcN that explicitly accounts for the energetic cost of its cryptic plasmids and achieves improved quantitative agreement with experimental data. Comparative analyses with previous models revealed that increased structural complexity does not necessarily enhance predictive accuracy, underscoring the importance of contextual and condition-specific validation. Metabolomic analyses further demonstrated that EcN exhibits a distinctive metabolic phenotype under gut-like conditions, characterized by enhanced amino acid consumption and short-chain fatty acid production, reinforcing its probiotic identity. Overall, this work provides a robust framework for the quantitative and physiologically relevant modeling of EcN and related commensal bacteria, with implications for probiotic research, industrial fermentation, and synthetic biology applications.

## Author Contributions

**Paola Corbín Agustí**: Conceptualization, Methodology, Formal analysis, Visualization, Investigation, Software, Data curation, Writing– Original draft, Reviewing and Editing. **Alba Arévalo-Lalanne:** Methodology, Investigation, Writing– Reviewing and editing. **Patricia Álvarez**: Methodology, Formal analysis, Investigation, Software, Data curation, Writing– Reviewing and editing. **María Enrique:** Methodology, Investigation, Writing– Reviewing and editing. **Daniel Ramón**: Conceptualization, Formal analysis, Writing– Reviewing and editing. **Juli Peretó**: Conceptualization, Formal analysis, Funding acquisition, Writing– Original draft, Reviewing and Editing. **Marta Tortajada:** Conceptualization, Formal analysis, Supervision, Writing– Original draft, Reviewing and Editing.

## Data Availability Statement

The code and data for reproducing this analysis are available in the GitHub repository: https://github.com/paocorbin/EcN-GEM

