## Supplementary Figures and Tables for "Flux modelling analysis reveals the metabolic impact of cryptic plasmids and environmental conditions in probiotic *Escherichia coli* Nissle 1917"

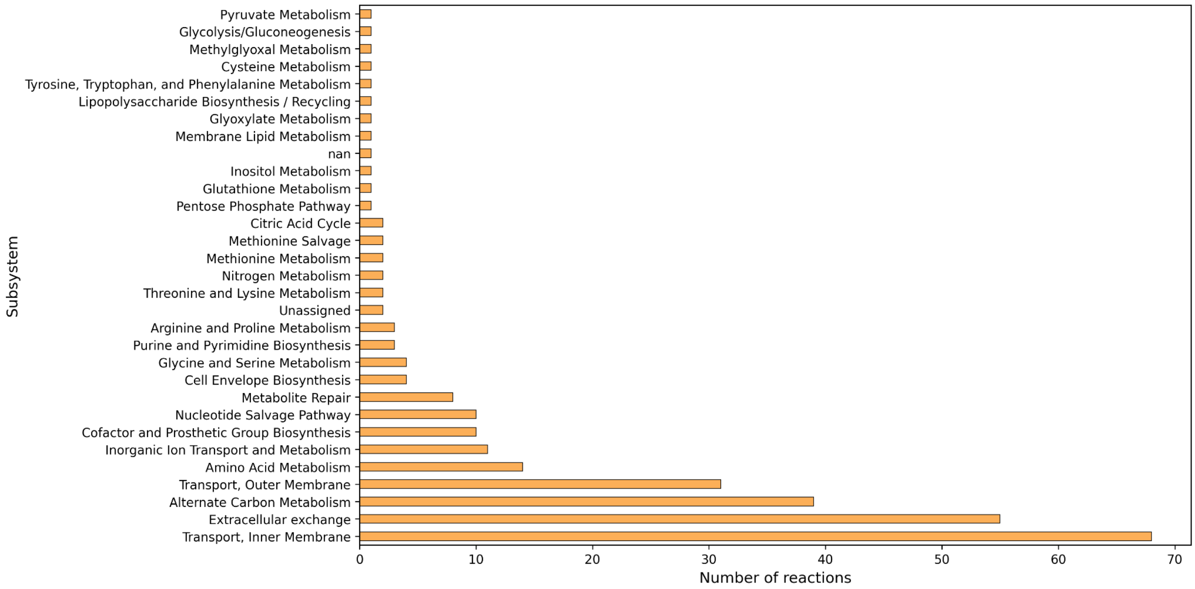


**Figure S1. Subsystems for reactions shared by models iHM1533 and iDK1463.**
Reactions common to both models were assigned to subsystems based on the reaction annotations in iHM1533.

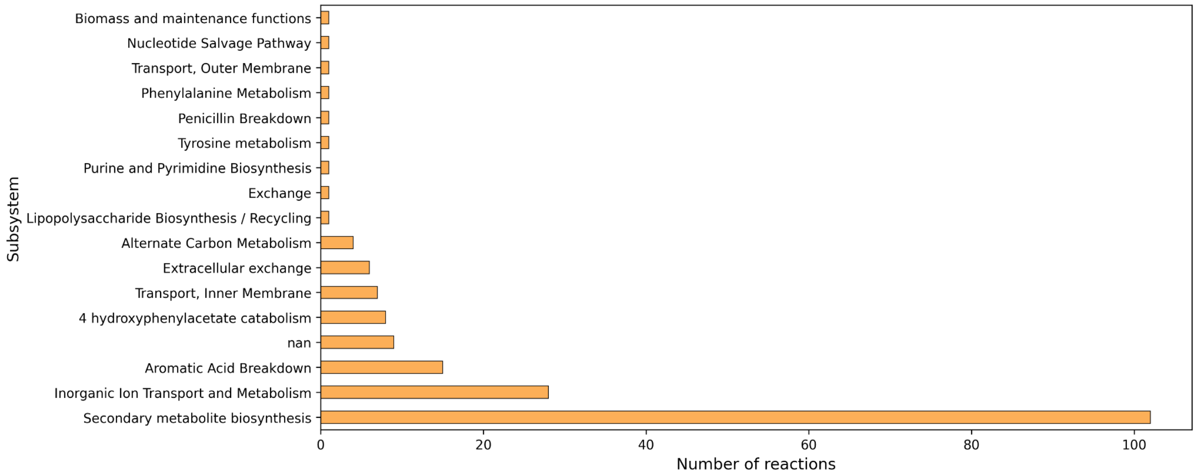


**Figure S2. Number of unique reactions per subsystem in iHM1533.**
Counts of reactions unique to iHM1533 (relative to the other EcN models) grouped by subsystem. Subsystems were assigned from the reaction annotations in iHM1533.

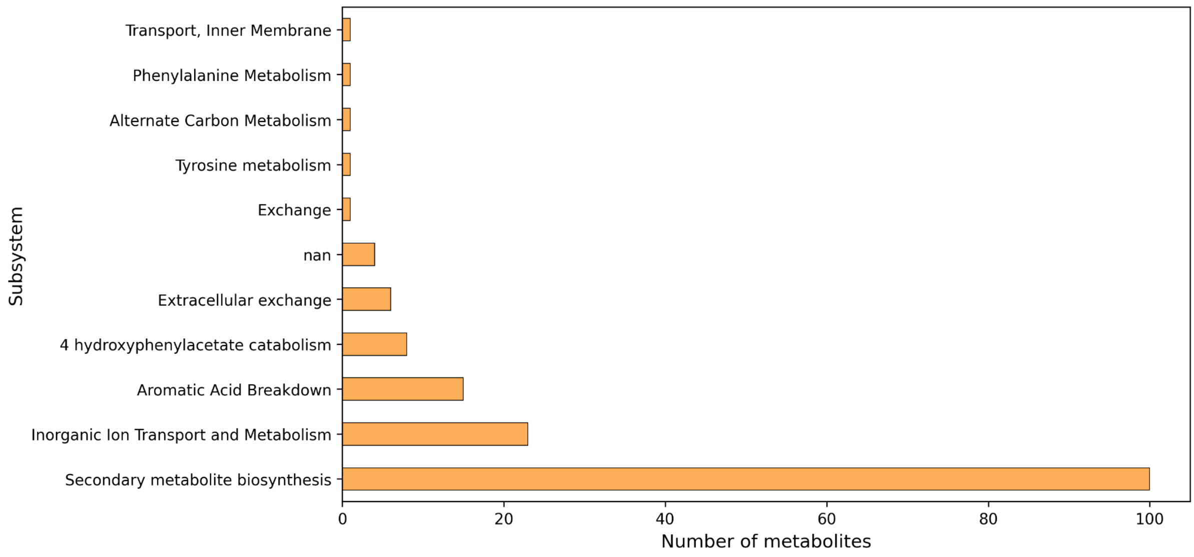


**Figure S3**. **Subsystems for unique metabolites in iHM1533.**
Subsystems were assigned from iHM1533 annotations and from the reactions associated with each metabolite.

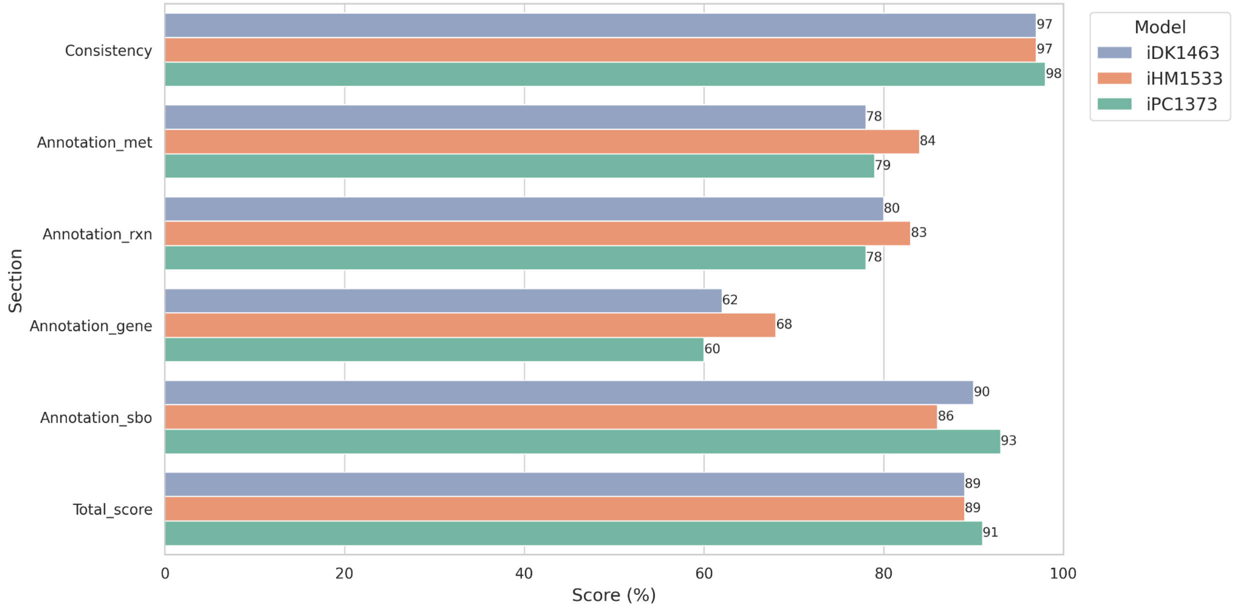


**Figure S4. Memote comparison of iDK1463, iHM1533 and iPC1373.**
The Memote test suite was used to compare metabolic network consistency and annotation coverage for metabolites (met), reactions (rxn), genes (gene) and SBO terms (sbo) in *E. coli* Nissle 1917 models. Scores for iDK1463 and iHM1533 models were taken from their respective publications.

**Table S1. Relative square errors (RSE) for the biomass growth rate predictions.**
RSE values were calculated by taking the squared difference between the predicted and experimental biomass growth rates and dividing it by the squared experimental value. For each model, the median RSE across all evaluated conditions was then obtained and expressed as a percentage.

| | **Models excluding plasmid consideration** | | |
|---|---|---|---|
| **Condition** | **iPC1373** | **iDK1463** | **iHM1533** |
| Glucose wt | 0.15 | 0.12 | 0.15 |
| Gluconate wt | 0.24 | 0.19 | 0.24 |
| Glucose ΔcsrA51 | 0.17 | 0.16 | 0.20 |
| Gluconate ΔcsrA51 | 0.46 | 0.39 | 0.47 |
| Fucose | 0.00 | 0.00 | 0.00 |
| Rhamnose | 0.04 | 0.04 | 0.06 |
| **Total median** | **16.3%** | **13.7%** | **17.7%** |
| | **Models including plasmid consideration** | | |
| **Condition** | **iPC1373** | **iDK1463** | **iHM1533** |
| Glucose wt | 0.02 | 0.01 | 0.02 |
| Gluconate wt | 0.06 | 0.04 | 0.06 |
| Glucose ΔcsrA51 | 0.09 | 0.08 | 0.11 |
| Gluconate ΔcsrA51 | 0.12 | 0.09 | 0.12 |
| Fucose | 0.04 | 0.04 | 0.02 |
| Rhamnose | 0.02 | 0.02 | 0.01 |
| **Total median** | **4.6%** | **3.6%** | **4.0%** |

**Table S2. Predicted external fluxes versus experimental measurements across different growth conditions.**

Metabolic fluxes for each model (iPC1373, iDK1463, and iHM533) are presented relative to each condition and dataset. The extracellular fluxes, both measured and predicted, include: substrate uptake ($q_S$), acetate formation ($q_{ace)}$, pyruvate formation ($q_{pyr}$), CO2 formation ($q_{CO2}$), glycogen formation ($q_{glyc}$) and 1,2-propanediol formation ($q_{12ppd}$). Underlined entries denote experimentally measured rates (Exp.) and those enforced as constraints during estimation. The extracellular flux results correspond to the FBA solution with the biomass reaction as the objective function.

| | | | $Q_s$ | $Q_{ace}$ | $Q_{CO2}$ | $Q_{pyr}$ | $Q_{glyc}$ | $Q_{12ppd}$ |
|---|---|---|---|---|---|---|---|---|
| **Condition** | **Reference** | **($h^{-1}$)** | **mmol·gDW$^{-1}$·h$^{-1}$** | | | | | |
| **Glc wt** | Exp. | <u>0.79</u> | <u>12.5 +- 1</u> | <u>6.3</u> | — | <u>n.d.</u> | <u>0.003</u> | — |
| | iPC1373 | 0.94 | <u>12.5 +- 1</u> | 10.05 | — | 0 | 0 | — |
| | iDK1463 | 0.91 | <u>12.5 +- 1</u> | 10.68 | — | 0 | 0 | — |
| | iHM1533 | 0.94 | <u>12.5 +- 1</u> | 10.67 | — | 0 | 0 | — |
| **Glc ΔcsrA51** | Exp. | <u>0.63</u> | <u>10.8 +- 0.7</u> | <u>7.2</u> | — | <u>n.d.</u> | <u>0.057</u> | — |
| | iPC1373 | 0.85 | <u>10.8 +- 0.7</u> | 5.87 | — | 0 | 0 | — |
| | iDK1463 | 0.83 | <u>10.8 +- 0.7</u> | 6.44 | — | 0 | 0 | — |
| | iHM1533 | 0.86 | <u>10.8 +- 0.7</u> | 6.44 | — | 0 | 0 | — |
| **Glcn wt** | Exp. | <u>0.74</u> | <u>20 +- 1.7</u> | <u>10.4</u> | — | <u>3.2</u> | <u>0.001</u> | — |
| | iPC1373 | 1.10 | <u>20 +- 1.7</u> | 19.65 | — | 0 | 0 | — |
| | iDK1463 | 1.06 | <u>20 +- 1.7</u> | 22.93 | — | 0 | 0 | — |
| | iHM1533 | 1.10 | <u>20 +- 1.7</u> | 23.12 | — | 0 | 0 | — |
| **Glcn ΔcsrA51** | Exp. | <u>0.57</u> | <u>12.6 +- 1.3</u> | <u>6.3</u> | — | <u>0.4</u> | <u>0.007</u> | — |
| | iPC1373 | 0.90 | <u>12.6 +- 1.3</u> | 8.65 | — | 0 | 0 | — |
| | iDK1463 | 0.87 | <u>12.6 +- 1.3</u> | 9.26 | — | 0 | 0 | — |
| | iHM1533 | 0.90 | <u>12.6 +- 1.3</u> | 9.28 | — | 0 | 0 | — |
| **Fucose** | Exp. | <u>0.48</u> | <u>12.8 +- 0.3</u> | <u>4</u> | <u>15.3</u> | — | — | <u>11.1</u> |
| | iPC1373 | 0.65 | <u>12.8 +- 0.3</u> | 12.03 | 14.76 | — | — | 0 |
| | iDK1463 | 0.71 | <u>12.8 +- 0.3</u> | 15.40 | 11.57 | — | — | 0 |
| | iHM1533 | 0.74 | <u>12.8 +- 0.3</u> | 15.29 | 11.22 | — | — | 0 |
| **Rhamnose** | Exp. | <u>0.28</u> | <u>6.9 +- 0.2</u> | <u>0.2</u> | <u>12.3</u> | — | — | <u>5.5</u> |
| | iPC1373 | 0.53 | <u>6.9 +- 0.2</u> | 1.33 | 17.52 | — | — | 0 |
| | iDK1463 | 0.57 | <u>6.9 +- 0.2</u> | 0.67 | 17.58 | — | — | 0 |
| | iHM1533 | 0.59 | <u>6.9 +- 0.2</u> | 0.64 | 17.45 | — | — | 0 |

For model rows (iPC1373, iDK1463 and iHM1533), Qs was constrained to the experimental value (Exp.); other fluxes are model predictions.

**Table S3. Relative square errors (RSE) for the external rates predictions.**
RSE values were calculated by taking the squared difference between the predicted and experimental flux rates and dividing it by the squared experimental value. For each model, the median RSE across all evaluated fluxes was then obtained and expressed as a percentage.

| Condition | Flux | iPC1373 | iDK1463 | iHM1533 |
|---|---|---|---|---|
| **Glucose wt** | $Q_{ace}$ | 0.35 | 0.48 | 0.48 |
| | $Q_{CO2}$ | — | — | — |
| | $Q_{pyr}$ | | | |
| | $Q_{glyc}$ | 0.00 | 0.00 | 0.00 |
| **Glucose ΔcsrA51** | $Q_{ace}$ | 0.03 | 0.01 | 0.01 |
| | $Q_{CO2}$ | — | — | — |
| | $Q_{pyr}$ | | | |
| | $Q_{glyc}$ | 0.00 | 0.00 | 0.00 |
| **Gluconate wt** | $Q_{ace}$ | 0.79 | 1.45 | 1.5 |
| | $Q_{CO2}$ | — | — | — |
| | $Q_{pyr}$ | 1.00 | 1.00 | 1.00 |
| | $Q_{glyc}$ | 1.00 | 1.00 | 1.00 |
| **Gluconate ΔcsrA51** | $Q_{ace}$ | 0.14 | 0.22 | 0.22 |
| | $Q_{CO2}$ | — | — | — |
| | $Q_{pyr}$ | 1.00 | 1.00 | 1.00 |
| | $Q_{glyc}$ | 0.00 | 0.00 | 0.00 |
| **Fucose** | $Q_{ace}$ | 4.03 | 8.12 | 7.97 |
| | $Q_{CO2}$ | 0 | 0.06 | 0.07 |
| | $Q_{pyr}$ | — | — | — |
| | $Q_{glyc}$ | — | — | — |
| **Rhamnose** | $Q_{ace}$ | 32.09 | 5.41 | 4.91 |
| | $Q_{CO2}$ | 0.18 | 0.18 | 0.18 |
| | $Q_{pyr}$ | — | — | — |
| | $Q_{glyc}$ | — | — | — |
| **Total median** | | **27%** | **35%** | **35%** |

**Table S4. Relative square errors (RSE) for central metabolism flux reactions.**
RSE values were calculated by taking the squared difference between the predicted and experimental flux rates and dividing it by the squared experimental value. For each model, the median RSE across all evaluated reaction fluxes, for each condition and methodology, was then obtained and expressed as a percentage. Due to clearance, only RSE median data is shown.

| Condition | Methodology | iPC1373 | iDK1463 | iHM1533 |
|---|---|---|---|---|
| **Glucose** | **FBA** | 26% | 27% | 27% |
| | **Possibilistic PFA** | 21% | 100% | 11% |
| **Gluconate** | **FBA** | 26% | 29% | 30% |
| | **Possibilistic PFA** | 100% | 51% | 100% |
| **Overall performance** | **FBA** | 31% | 29% | 33% |
| | **Possibilistic PFA** | 32% | 100% | 50% |

**Table S5. Mapping of central-carbon metabolite conversions to reactions in metabolic models.**
This table lists, for each reaction index in the figure, the metabolite conversion (Revelles et al., 2013) and the corresponding representative reactions in metabolic models. Reaction IDs follow BiGG nomenclature; reversible steps are indicated with "↔", a leading "−" in reaction ID denotes use of the reverse direction, and exchange reactions are labeled EX_.

| Reaction Index | Metabolite conversion (Revelles reactions) | Model reactions IDs |
|---|---|---|
| **1** | Glucose + PEP → G6P + Pyr / Gluconate → 6PG | GLCptspp / GNK, -GNP |
| **2** | F6P ↔ G6P | PGI |
| **3** | F6P ↔ FBP | PFK, -FBP |
| **4** | FBP ↔ GAP + GAP | FBA, DKFPS2, TPI |
| **5** | GAP ↔ PGA | PGM |
| **6** | PGA ↔ PEP | ENO |
| **7** | PEP ↔ Pyr | PYK, -PPS, TREptspp, PYK4, GLCptspp, MNLptspp, GAMptspp, SRBptspp, SUCptspp, ACGALptspp, CHITOBpts, GALTptspp, PYK2, FRUpts2pp, ACMANAptspp, SBTptspp, 2DGLCptspp, ASCBptspp, CELBpts, MALTptspp, |

| Reaction Index | Metabolite conversion (Revelles reactions) | Model reactions IDs |
|---|---|---|
| | | DHAPT, MANptspp, MANGLYCptspp, CELLBpts, CHTBSptspp, PPS, GALAMPTSpp, ARBTptspp, PYK3, ACGAptspp, CLBptspp, ACMUMptspp, PYK6, FRUptspp, SALCptspp |
| **8** | OAA + AcCoA → Cit | CS |
| **9** | Cit → AKG + CO2 | ICDHyr |
| **10** | AKG → Suc + CO2 | AKGDH, AKGDH2, TAUDO |
| **11** | Suc → Mal | SUCDi, FUM |
| **12** | Mal ↔ OAA | MDH3, MOX, MDH2, MDH, DMALRED |
| **13** | PEP + CO2 ↔ OAA | PPC, -PPCK |
| **14** | Pyr → AcCoA + CO2 | PFOR, POR5, PDH, PFL |
| **15** | Mal → Pyr + CO2 | ME1, ME2 |
| **16** | 6PG → GAP + Pyr | EDA |
| **17** | G6P → 6PG | G6PDH2r, PGL |
| **18** | 6PG → R5P + CO2 | RPI |
| **19** | R5P ↔ GAP + E2 | TKT1 |
| **20** | F6P ↔ E4P + E2 | TKT2 |
| **21** | R5P + E2 ↔ S7P | TKT1 |
| **22** | GAP + E3 ↔ F6P | TALA |
| **23** | AcCoA → Acetate | ACOXT, ACACCT, -ACS, HXCT, BUTCT, -ACKr |
| **24** | Pyr → Pyruvate | EX_pyr_e |
