## Supplementary Methods for "Flux modelling analysis reveals the metabolic impact of cryptic plasmids and environmental conditions in probiotic *Escherichia coli* Nissle 1917"

### 1. Bacterial culture conditions

Experiments utilized M9 minimal medium supplemented with glucose (Table SM1) and rich-gut microbiota medium (GMM) (Table SM2).

**Table SM1. Composition of the M9 minimal medium with glucose.** The medium was prepared following the methodology described in Monk et al. (2016). Additional details regarding its preparation can be found in the STAR Methods section of the aforementioned publication.

| **Component** | **Concentration** |
|---|---|
| M9 Minimal Medium Salts 5X | 1X |
| $MgSO_4$ | 2mM |
| $CaCl_2$ | 0.1mM |
| ATCC Vitamin Mix | 1% |
| ATCC Trace Mineral Mix | 1% |
| Thiamine | 1mM |
| Casamino acids | 0.2% |
| D-Glucose | 2g/L |

**Table SM2. Composition of the Rich Gut Microbiota Medium (GMM)**. The medium's composition has been previously detailed by Goodman et al. (2011) and Średnicka et al. (2023), with the latter describing it in an agar-free formulation. Further information regarding this medium can be found within the methodology sections of both articles.

| Component | Concentration |
|---|---|
| Tryptone peptone | 0.2% |
| Yeast extract | 0.1% |
| D-Glucose | 2.2mM |
| L-Cysteine | 3.2mM |
| Cellobiose | 2.9mM |
| Maltose | 2.8mM |
| Fructose | 2.2mM |
| Meat extract | 0.5% |
| $KH_2PO_4$ | 100mM |
| $MgSO_4$-$7H_2O$ | 0.008mM |
| $NaHCO_3$ | 4.8mM |
| $NaCl_2$ | 1.37mM |
| $CaCl_2$ | 0.80% |
| Vitamin K (Menadione) | 5.8mM |
| $FeSO_4$ | 1.44mM |
| Tween 80 | 0.05% |
| ATCC Vitamin Mix | 1% |
| ATCC Trace Mineral Mix | 1% |
| Acetic acid | 30mM |
| Isovaleric acid | 1mM |
| Propionic acid | 8mM |
| Butyric acid | 4mM |
| pH adjusted to 6.5 | |

### 2. ATP expenditure parameters: NGAM

To establish the Non-Growth Associated Maintenance Energy (NGAM) and the bounds of the ATPM reaction, while accounting for the metabolic burden linked to plasmid maintenance, we used the median of previously reported NGAM values (Table SM3). This median was applied as the lower bound of the ATPM reaction under aerobic (25 mmol/gDW·h) and anaerobic (3.5 mmol/gDW·h) conditions.

**Table SM3. Non-Growth Associated Maintenance Energy (NGAM) Values Plasmid-Bearing *Escherichia coli*.** This table compiles non-growth associated maintenance energy (NGAM) values, expressed in mmol·gDW$^{-1}$·h$^{-1}$, for plasmid-bearing *E. coli*. Data presented in this table are the corresponding article reference, the specific organism studied, the oxygen condition, and the NGAM value.

| Reference | Organism | Condition | NGAM (mmol/gDW·h) |
|---|---|---|---|
| Oftadeh & Hatzimanikatis, 2024 | *E. coli* containing pMB1 | Aerobe | 30 |
| | *E. coli* containing Ori2 | Aerobe | 15 |
| Zeng & Yang, 2019 | *E. coli* W3110 GFP-bearing | Aerobe | 32.6 |
| Weber et al., 2002 | *E. coli* GMO, Induced | Aerobe | 20 |
| Boecker et al., 2021 | *E. coli MG1655* pLC control | Anaerobe | 3.30 ± 0.43 |
| | *E. coli MG1655* pMC control | Anaerobe | 3.61 ± 0.82 |
| | *E. coli MG1655* pHC control | Anaerobe | 2.67 ± 1.05 |

### 3. Plasmid integration in the EcN model and sensitivity analysis

To simulate plasmid replication and its associated metabolic burden, a fixed lower bound was imposed on the plasmid reaction while maintaining biomass maximization as the objective function. The appropriate lower bound was determined by evaluating a range of NGAM values (1–50 mmol·gDW⁻¹·h⁻¹) to identify the configuration that best reproduced the experimental conditions (Table SM6) and to assess model sensitivity to NGAM variation. As a control, models incorporating the plasmid reaction but without any additional constraint on this flux were initially simulated under aerobic conditions using four carbon sources (glucose, gluconate, fucose, and rhamnose), with biomass maximization as the objective. Subsequently, lower bounds ranging from 0.1 to 1.4 mmol·gDW⁻¹·h⁻¹ were applied to the plasmid reaction. All simulations were constrained using the available experimental data for each condition (Table SM6), specifically by restricting carbon source uptake rates and secretion or by-product fluxes within their experimental range (mean ± SD). Given the central role of acetate overflow in representing metabolic burden, an additional set of simulations was performed with the acetate flux left unconstrained to examine its behavior. The outcomes of these analyses are summarized in Tables SM4 and SM5.

**Table SM4. Simulation of NGAM values under different plasmid and acetate constraints.** Non-growth-associated maintenance energy (NGAM, mmol·gDW⁻¹·h⁻¹) predicted from FBA simulations optimizing biomass reaction for the EcN models iDK1463, iHM1533, and iPC1373. Simulations were performed under fixed plasmid reaction lower bounds (0.0 to 1.4 mmol·gDW⁻¹·h⁻¹), with the acetate exchange reaction either constrained to the experimental value (+) or left unconstrained (-). The NGAM value for each model shown corresponds to the mean at which the simulated biomass flux falls within the experimental growth range. The “Mean” column represents the average NGAM across the three models for each condition.

| | | | **NGAM result** (mmol·gDW⁻¹·h⁻¹) | | | |
|---|---|---|---|---|---|---|
| **Carbon source** | **Plasmid rxn lower bound** | **Acetate constrained** | **iDK1463** | **iHM1533** | **iPC1373** | **Mean** |
| **Glucose** | **0** | **+** | 33 | 35.5 | 35.5 | 34.67 |
| | | **-** | 39 | 41.5 | 41.5 | 40.67 |
| **Gluconate** | | **+** | 39.5 | 42 | 41.5 | 41.00 |
| | | **-** | 51.5 | 54 | 54 | 53.17 |
| **Rhamnose** | | **+** | 19 | 21 | 19 | 19.67 |
| | | **-** | 19.5 | 21 | 19 | 19.83 |
| **Fucose** | | **+** | 10.5 | 14 | 10.5 | 11.67 |
| | | **-** | 21 | 24 | 21 | 22.00 |
| **Glucose** | **0.5** | **+** | 32.50 | 35.50 | 35.50 | 34.50 |

| | | | **NGAM result** (mmol·gDW⁻¹·h⁻¹) | | | |
|---|---|---|---|---|---|---|
| **Carbon source** | **Plasmid rxn lower bound** | **Acetate constrained** | **iDK1463** | **iHM1533** | **iPC1373** | **Mean** |
| | | **-** | 38.50 | 41.50 | 41.50 | 40.50 |
| **Gluconate** | | **+** | 38.50 | 41.50 | 41.50 | 40.50 |
| | | **-** | 51.50 | 53.50 | 53.50 | 52.83 |
| **Rhamnose** | | **+** | 18.50 | 20.00 | 18.00 | 18.83 |
| | | **-** | 18.50 | 20.50 | 18.50 | 19.17 |
| **Fucose** | | **+** | 10.00 | 13.00 | 10.00 | 11.00 |
| | | **-** | 20.00 | 23.50 | 20.00 | 21.17 |
| **Glucose** | **1** | **+** | 32.50 | 35.00 | 35.00 | 34.17 |
| | | **-** | 38.00 | 41.00 | 40.50 | 39.83 |
| **Gluconate** | | **+** | 38.50 | 41.00 | 41.00 | 40.17 |
| | | **-** | 50.50 | 53.50 | 53.50 | 52.50 |
| **Rhamnose** | | **+** | 17.50 | 19.50 | 17.50 | 18.17 |
| | | **-** | 18.00 | 20.00 | 17.50 | 18.50 |
| **Fucose** | | **+** | 9.50 | 12.50 | 9.50 | 10.50 |
| | | **-** | 19.50 | 23.00 | 19.50 | 20.67 |
| **Glucose** | **1.4** | **+** | 32.00 | 34.50 | 34.50 | 33.67 |
| | | **-** | 37.50 | 40.50 | 40.50 | 39.50 |
| **Gluconate** | | **+** | 38.50 | 40.50 | 40.50 | 39.83 |
| | | **-** | 50.50 | 53.50 | 53.50 | 52.50 |
| **Rhamnose** | | **+** | 17.00 | 19.00 | 17.00 | 17.67 |
| | | **-** | 17.50 | 19.50 | 17.00 | 18.00 |
| **Fucose** | | **+** | 8.50 | 12.00 | 8.50 | 9.67 |
| | | **-** | 19.00 | 22.00 | 19.00 | 20.00 |

**Table SM5. Mean and median NGAM values from FBA simulations under different plasmid constraints.** Average and median non-growth-associated maintenance energy (NGAM, mmol·gDW$^{-1}$·h$^{-1}$) obtained from FBA simulations optimizing biomass across all EcN models and carbon sources in Table SM3.

| **Plasmid rxn lower bound** | **NGAM result** (mmol·gDW$^{-1}$·h$^{-1}$) | |
|---|---|---|
| | **Mean** | **Median** |
| 0 | 26.75 | 27.17 |
| 0.5 | 26.21 | 26.67 |
| 1 | 25.75 | 26.17 |
| 1.4 | 25.21 | 25.67 |

The resulting NGAM values exhibited substantial variation across carbon sources to match the experimental biomass production rates (Table SM4). When the acetate flux was left unconstrained, the mean NGAM value increased consistently across all simulations. This result underscores the importance of appropriately constraining acetate exchange to accurately reproduce the experimental flux distribution under specific growth conditions. Although no major discrepancies were observed among the three models, all required higher NGAM fluxes when acetate was unconstrained, suggesting that none of the models could fully capture the experimentally observed acetate flux. The NGAM sensitivity analysis further highlighted the critical influence of properly defining the flux through the ATPM reaction.

Overall, a lower bound of 1 mmol·gDW$^{-1}$·h$^{-1}$ was selected for the plasmid reaction, as this configuration yielded NGAM values closely aligned with the reported average of 25 mmol·gDW$^{-1}$·h$^{-1}$ (Tables SM3 and SM5).

### 4. Experimental datasets employed for model validation

Experimental data from the literature on EcN growth across multiple carbon sources were compiled for model validation (Revelles et al., 2013; Millard et al., 2021). These datasets contain experimental extracellular fluxes, including specific growth rate, substrate uptake rate, and secretion rates of key byproducts (Table SM6).

**Table SM6. Experimental datasets for extracellular fluxes.** The data shown are the specific growth rate ($h^{-1}$ substrate uptake ($q_S$), acetate formation ($q_{ace}$), pyruvate formation ($q_{pyr}$), $CO_2$ formation ($q_{CO2}$), glycogen formation ($q_{glyc}$) and 1,2-propanediol formation ($q_{12ppd}$) compiled from Revelles et al. (2013)* and Millard et al. (2021)** in aerobic conditions.

| | | $Q_s$ | $Q_{ace}$ | $Q_{CO2}$ | $Q_{pyr}$ | $Q_{glyc}$ | $Q_{12ppd}$ |
|---|---|---|---|---|---|---|---|
| **Condition** | **μ ($h^{-1}$)** | **mmol·gDW⁻¹·h⁻¹** | | | | | |
| **Glucose wild-type*** | 0.79 ± 0.02 | 12.5 ± 1.0 | 6.3 ± 0.1 | — | n.d. | 0.003 ± 0.001 | — |
| **Glucose Δ*csrA*51*** | 0.63 ± 0.02 | 10.8 ± 0.7 | 7.2 ± 0.1 | — | n.d. | 0.057 ± 0.014 | — |
| **Gluconate wild-type*** | 0.74 ± 0.02 | 20.0 ± 1.7 | 10.4 ± 0.1 | — | 3.2 ± 0.1 | 0.001 ± 0.001 | — |
| **Gluconate Δ*csrA*51*** | 0.57 ± 0.01 | 12.6 ± 1.3 | 6.3 ± 0.1 | — | 0.4 ± 0.3 | 0.007 ± 0.001 | — |
| **Fucose**** | 0.48 ± 0.01 | 12.8 ± 0.3 | 4.0 ± 0.2 | 15.3 ± 2.4 | — | — | 11.1 ± 0.2 |
| **Rhamnose**** | 0.28 ± 0.01 | 6.9 ± 0.2 | 0.2 ± 0.1 | 12.3 ± 0.7 | — | — | 5.5 ± 0.1 |

*n.d.*, not detected; —, not available.

### 5. *In silico* constraints and growth conditions

For reversible reactions, upper and lower bounds of –1000 and 1000 mmol·gDW⁻¹·h⁻¹, respectively, were applied. For irreversible reactions, the lower bound –or in specific cases, the upper bound– was fixed at 0 mmol·gDW⁻¹·h⁻¹. Maximization of growth rate was set as the objective function.

Anaerobic conditions were imposed by constraining the lower bound of the $O_2$ exchange reaction to 0 mmol·gDW⁻¹·h⁻¹; whereas aerobic conditions were simulated by setting this bound to -20 mmol·gDW⁻¹·h⁻¹. The growth-associated maintenance energy (GAM) was fixed at 59.8 mmol·gDW⁻¹·h⁻¹, while the non-growth-associated maintenance energy (NGAM) was set to 25 mmol·gDW⁻¹·h⁻¹ under aerobic and 3.5 mmol·gDW⁻¹·h⁻¹ under anaerobic conditions. When plasmid maintenance was included, the lower bound of the rxn_plasmid_c reaction was set to 1 mmol·gDW⁻¹·h⁻¹.

For the simulation of each growth media, the lower bounds of all exchange reactions were set to zero except for the metabolites present in the defined medium: M9 minimal medium (Table SM7) or gut microbiota medium GMM (Table SM8).

**Table SM7. Definition of *in silico* M9 glucose minimal medium.** The lower and upper bounds (mmol·gDW$^{-1}$·h$^{-1}$) of the exchange reactions that were constrained to simulate the M9 medium.

| Reaction ID | Met Name | Lower bound (mmol·gDW$^{-1}$·h$^{-1}$) | Upper bound (mmol·gDW$^{-1}$·h$^{-1}$) |
|---|---|---|---|
| EX_co2_e | CO2 | -1000 | 1000 |
| EX_h2o_e | H2O | -1000 | 1000 |
| EX_h_e | H | -1000 | 1000 |
| EX_ca2_e | Calcium | -1000 | 1000 |
| EX_cl_e | Chloride | -1000 | 1000 |
| EX_cobalt2_e | Co2 | -1000 | 1000 |
| EX_cu2_e | Cu2 | -1000 | 1000 |
| EX_fe2_e | Iron (Fe2+) | -1000 | 1000 |
| EX_fe3_e | Iron (Fe3+) | -1000 | 1000 |
| EX_k_e | Potassium | -1000 | 1000 |
| EX_mg2_e | Mg | -1000 | 1000 |
| EX_mn2_e | Mn2 | -1000 | 1000 |
| EX_mobd_e | Molybdate | -1000 | 1000 |
| EX_na1_e | Sodium | -1000 | 1000 |
| EX_nh4_e | Ammonia | -1000 | 1000 |
| EX_ni2_e | Ni2 | -1000 | 1000 |
| EX_pi_e | Phosphate | -1000 | 1000 |
| EX_sel_e | Selenate | -1000 | 1000 |
| EX_so4_e | Sulfate | -1000 | 1000 |
| EX_zn2_e | Zinc | -1000 | 1000 |
| EX_cbl1_e* | Cobalamin (I) | -0.01 | 1000 |
| EX_thm_e** | Thiamin | -0.01 | 1000 |

| Reaction ID | Met Name | Lower bound (mmol·gDW⁻¹·h⁻¹) | Upper bound (mmol·gDW⁻¹·h⁻¹) |
|---|---|---|---|
| EX_glc__D_e*** | D-Glucose | -20 | 1000 |

*Monk et al. (2016); **Nicolas et al. (2007); ***lower bound of D-Glucose was set to -20 mmol·gDW⁻¹·h⁻¹ just in the metabolomics data simulation. In other simulations, the carbon source was changed according to its simulation particularities.

**Table SM8. Definition of *in silico* Gut Microbiota Medium (GMM).** The lower and upper bounds of the exchange reactions that were constrained to simulate the GMM medium. This composition was modified from van't Hof et al., 2022.

| Reaction ID | Met Name | Lower bound (mmol·gDW⁻¹·h⁻¹) | Upper bound (mmol·gDW⁻¹·h⁻¹) |
|---|---|---|---|
| EX_3mb_e | 3-Methylbutanoic acid | -1000 | 1000 |
| EX_4abz_e | 4-Aminobenzoate | -1000 | 1000 |
| EX_ac_e | Acetate | -1000 | 1000 |
| EX_btn_e | Biotin | -1000 | 1000 |
| EX_but_e | Butyrate | -1000 | 1000 |
| EX_ca2_e | Calcium | -1000 | 1000 |
| EX_cbl1_e | Cobalamin (I) | -1000 | 1000 |
| EX_cbl2_e | Cobalamin (II) | -1000 | 1000 |
| EX_cellb_e | Cellobiose | -1000 | 1000 |
| EX_cl_e | Chloride | -1000 | 1000 |
| EX_cobalt2_e | Cobalt (Co2+) | -1000 | 1000 |
| EX_cu2_e | Copper | -1000 | 1000 |
| EX_cys__L_e | L-Cysteine | -1000 | 1000 |
| EX_fe2_e | Iron (Fe2+) | -1000 | 1000 |
| EX_fe3_e | Iron (Fe3+) | -1000 | 1000 |
| EX_fol_e | Folate | -1000 | 1000 |
| EX_fru_e | D-Fructose | -1000 | 1000 |

| Reaction ID | Met Name | Lower bound (mmol·gDW⁻¹·h⁻¹) | Upper bound (mmol·gDW⁻¹·h⁻¹) |
|---|---|---|---|
| EX_glc__D_e | D-Glucose | -1000 | 1000 |
| EX_h_e | H+ | -1000 | 1000 |
| EX_h2o_e | H2O | -1000 | 1000 |
| EX_hco3_e | Bicarbonate | -1000 | 1000 |
| EX_his__L_e | L-Histidine | -1000 | 1000 |
| EX_k_e | Potassium | -1000 | 1000 |
| EX_lipoate_e | Lipoate | -1000 | 1000 |
| EX_malt_e | Maltose | -1000 | 1000 |
| EX_mg2_e | Magnesium | -1000 | 1000 |
| EX_mn2_e | Manganese | -1000 | 1000 |
| EX_mndn_e | Menadione | -1000 | 1000 |
| EX_mobd_e | Molybdate | -1000 | 1000 |
| EX_na1_e | Sodium | -1000 | 1000 |
| EX_nac_e | Nicotinate | -1000 | 1000 |
| EX_ni2_e | Nickel | -1000 | 1000 |
| EX_no3_e | Nitrate | -1000 | 1000 |
| EX_pheme_e | Protoheme | -1000 | 1000 |
| EX_pi_e | Phosphate | -1000 | 1000 |
| EX_pnto__R_e | (R)-Pantothenate | -1000 | 1000 |
| EX_ppa_e | Propionate | -1000 | 1000 |
| EX_pydxn_e | Pyridoxine | -1000 | 1000 |
| EX_ribflv_e | Riboflavin | -1000 | 1000 |
| EX_slnt_e | Selenite | -1000 | 1000 |
| EX_so4_e | Sulfate | -1000 | 1000 |

| Reaction ID | Met Name | Lower bound (mmol·gDW$^{-1}$·h$^{-1}$) | Upper bound (mmol·gDW$^{-1}$·h$^{-1}$) |
|---|---|---|---|
| EX_thm_e | Thiamin | -1000 | 1000 |
| EX_tungs_e | Tungstate | -1000 | 1000 |
| EX_zn2_e | Zinc | -1000 | 1000 |
| EX_ala__L_e | L-Alanine | -1000 | 1000 |
| EX_asn__L_e | L-Asparagine | -1000 | 1000 |
| EX_asp__L_e | L-Aspartate | -1000 | 1000 |
| EX_glu__L_e | L-Glutamate | -1000 | 1000 |
| EX_gln__L_e | L-Glutamine | -1000 | 1000 |
| EX_gly_e | Glycine | -1000 | 1000 |
| EX_ile__L_e | L-Isoleucine | -1000 | 1000 |
| EX_leu__L_e | L-Leucine | -1000 | 1000 |
| EX_lys__L_e | L-Lysine | -1000 | 1000 |
| EX_phe__L_e | L-Phenylalanine | -1000 | 1000 |
| EX_pro__L_e | L-Proline | -1000 | 1000 |
| EX_ser__L_e | L-Serine | -1000 | 1000 |
| EX_thr__L_e | L-Threonine | -1000 | 1000 |
| EX_trp__L_e | Tryptophan | -1000 | 1000 |
| EX_tyr__L_e | L-Tyrosine | -1000 | 1000 |
| EX_val__L_e | L-Valine | -1000 | 1000 |
| EX_arg__L_e | L-Arginine | -1000 | 1000 |
| EX_met__L_e | L-Methionine | -1000 | 1000 |

M9 minimal media was employed when simulating the experimental datasets, with additional condition-specific adjustments applied as required.

**Prediction of external rates: biomass growth rate**

To predict biomass growth rates under various conditions (Table SM6), the biomass reaction was defined as the objective function, and flux balance analysis (FBA) was performed to estimate the corresponding flux. For each condition, all experimentally measured fluxes were constrained within their reported ranges, except for biomass, which was left unconstrained to be predicted. Simulations were conducted using M9 minimal medium (Table SM7), varying only the carbon source according to each experimental dataset. Specifically, carbon source uptake and secretion or by-product reactions were restricted to bounds corresponding to the experimental range (mean ± SD). The aerobic condition was simulated by setting the lower bound of the $O_2$ exchange reaction to –20 mmol·gDW$^{-1}$·h$^{-1}$.

For simulations performed without plasmid consideration, the lower bound of the ATP maintenance (ATPM) reaction (NGAM) was fixed at 8.4 mmol·gDW$^{-1}$·h$^{-1}$. When plasmid maintenance was included, NGAM was adjusted as previously described for aerobic conditions, and the lower bound of the rxn_plasmid_c reaction was fixed at 1 mmol·gDW$^{-1}$·h$^{-1}$. To reproduce the experimental conditions reported by Revelles et al. (2013) for the wild-type and ΔcsrA51 mutant strains, additional constraints were introduced to account for the regulation of glycogen metabolism. Specifically, glycogen branching reactions (GLBRAN2, GLDBRAN2, and GLCP) were blocked by setting both their lower and upper bounds to zero. Furthermore, a sink reaction for glycogen (SK_glycogen_c) was added to simulate glycogen accumulation, following the approach of Morin et al. (2016).

**Prediction of external rates: remaining external rates**

To predict the external fluxes for which experimental data were available, we performed flux balance analysis (FBA), metabolic flux analysis (MFA; consistency = 1), and flux variability analysis (FVA; 90 % of the optimal objective value). In these simulations, only models including plasmid maintenance were used, applying the same constraints described previously for biomass growth rate simulations. However, in this case, only the substrate uptake rate for each condition (Table SM6) was constrained within its experimental range. All other measured fluxes were treated as variables to be predicted.

**Prediction of internal rates**

In this case, intracellular flux data were available only for the conditions reported by Revelles et al. (2013) (Table SM6), namely glucose and gluconate as carbon sources. Flux distributions were simulated using both MFA (consistency = 1) and FBA with biomass as the objective function. These simulations were conducted under aerobic conditions, using M9 minimal medium and including plasmid maintenance, applying the same constraints as those used for the prediction of biomass growth rate.

The metabolite conversions analyzed and their associated reactions are listed in Table S2. For each metabolite conversion, a net flux was calculated when multiple reactions could contribute to the same transformation. For the following reactions: GNP, FBP, PPCK, ACKr, ACS, PPS, and TKT2, the flux direction was inverted to ensure consistent alignment for net flux computation. In the case of conversions 11 and 17 (Table S2), the reaction with the lowest absolute flux was used to represent the transformation. Finally, all net fluxes were normalized to the respective glucose or gluconate uptake rate, setting this value to 100. These net and normalized fluxes were used to represent the predicted intracellular flux distributions for each model and analysis (FBA or MFA).

**Prediction of metabolomics data**

To predict the metabolites consumed and produced under anaerobic conditions in M9 medium with glucose and in GMM medium, the exchange reactions of the models were constrained according to Tables SM7 and SM8, respectively. Only the models including plasmid reactions were used, with NGAM set to 3.5 mmol·gDW$^{-1}$·h$^{-1}$ and the lower bound of the rxn_plasmid_c reaction fixed at 1 mmol·gDW$^{-1}$·h$^{-1}$. In these simulations, the bounds of the BUTCT reaction were adjusted to (–1000, 1000) to allow butyrate production under anaerobic conditions.
Flux distributions were computed using FBA, with biomass maximization as the objective function. Only the consumed and produced metabolites for which experimental data are available are shown (Supplementary Material 7), to evaluate the predictive performance of the models.

### 6. Metabolomic assays

The metabolites enumerated in Table SM5 were quantified utilizing high-performance liquid chromatography (HPLC). In this study various advanced analytical techniques were used with three different detection methods (HPLC-RID, HPLC-FLD, and LC-MS (single quadrupole mass spectrometer)), to identify and quantify potentially relevant metabolites in probiotic function. Metabolite selection was based on scientific literature, detailed in Table SM9.

**Table SM9. List of compounds for HPLC analysis of EcN.** Metabolites analyzed in EcN using high-performance liquid chromatography (HPLC). The table classifies the metabolites into various categories: short-chain fatty acids (SCFAs) and fermentation-related metabolites, arginine deiminase pathway metabolites, indole-based metabolites, and GABA metabolites. For each compound, its name, the analytical technique used for its quantification, and its exchange reaction BIGG ID (if available) are provided.

| **Compounds** | **Name** | **Analytical technique** | **BIGG ID exchange reaction** |
|---|---|---|---|
| **Short chain fatty acids (SCFAs) and fermentation-related metabolites** (Amrofell & Moon, 2023; Monk et al., 2016) | | | |
| Acetate | Acetic acid (C2:0) | HPLC-RID | EX_ac_e |
| Butyrate | Butyric acid (C4:0) | HPLC-RID | EX_but_e |
| Ethanol | Ethanol | HPLC-RID | EX_etoh_e |
| Formate | Formic acid (C1:0) | HPLC-RID | EX_for_e |
| Isobutyrate | Isobutyric acid (C4:0) | HPLC-RID | EX_ibt_e or EX_isobuta_e |
| Isovalerate | Isovaleric acid (C5:0) | HPLC-RID | EX_ival_e |
| Lactate | Lactic acid | HPLC-RID | EX_lac__D_e |
| Propionate | Propionic acid (C3:0) | HPLC-RID | EX_ppa_e |

| **Compounds** | **Name** | **Analytical technique** | **BIGG ID exchange reaction** |
|---|---|---|---|
| Succinate | Succinic acid | HPLC-RID | EX_succ_e |
| Valerate | Valeric acid (C5:0) | HPLC-RID | EX_pta_e |
| **Arginine deiminase pathway** (Van Der Hooft et al., 2019) | | | |
| Arginine | L-Arginine | HPLC-FLD | EX_arg__L_e |
| Ornithine | L-Ornithine | HPLC-FLD | EX_orn_e |
| **Indole-based metabolites** (Berstad et al., 2015; Michael et al., 2022) | | | |
| Tryptophan | L-Tryptophan | LC-MS | EX_trp__L_e |
| **GABA metabolites** (Mazzoli & Pessione, 2016; Pérez-Berezo et al., 2017) | | | |
| GABA | γ-Aminobutyric acid | LC-MS | EX_4abut_e |
| Glutamate | L-Glutamic acid | HPLC-FLD | EX_glu__L_e |
